# MYPT1 is a non-canonical AKAP that tethers PKA to the MLCP signaling node

**DOI:** 10.1101/2023.04.27.538407

**Authors:** Jawad S Khalil, Paulo A. Saldanha, Connor M Blair, Jiayue Ling, Wei Ji, George S. Baillie, Khalid M Naseem, Leonid L Nikitenko, Francisco Rivero

## Abstract

The activity of myosin light chain phosphatase (MLCP) is fine-tuned by the phosphorylation status of the MLCP target subunit 1 (MYPT1), which is determined by the antagonistic effects of Rho kinase (ROCK) and cAMP/cGMP-dependent protein kinases (PKA and PKG). PKA is composed of two regulatory (PKA-R, of which four variants exist) and two catalytic (PKAcat) subunits. PKA is targeted to the vicinity of its substrates by binding to A kinase anchoring proteins (AKAPs). MYPT1 is part of a complex signaling node that includes kinases and other enzymes involved in signal transduction. We hypothesized that MYPT1 might function as an AKAP to target PKA to the MLCP signaling node. Using a combination of immunoprecipitation, affinity pulldown and *in situ* proximity ligation assay (PLA) in human platelets and endothelial cells, we show that MYPT1 directly interacts with all four PKA-R variants and mapped the interaction to a 200 residues long central region of MYPT1. The interaction does not involve the docking and dimerization domain of PKA-R typically required for binding to AKAPs. Using peptide array overlay we identified K595, E676 and the PKA/ROCK kinase substrate motif R693/R694/S695/T696 as critical for the interaction. Substitution of S695, T696 or both by aspartic acid or the corresponding phosphorylated residue abolished binding. Our findings reveal that MYPT1 functions as a non-canonical AKAP to anchor PKA to the vicinity of non-phosphorylated S695/T696, where PKA-R would prevent PKAcat, and potentially also ROCK, from interacting with and phosphorylating MYPT1.

## Introduction

Regulation of actomyosin-driven contraction is critical for numerous cell functions in the cardiovascular system, including smooth muscle tone, changes in permeability of endothelial cell monolayers and platelet shape change and spreading. The principal regulatory mechanism of these processes involves the phosphorylation and dephosphorylation of the 20-kDa myosin II regulatory light chains (MLC), catalyzed by myosin light chain kinase (MLCK) and myosin light chain phosphatase (MLCP) respectively (1)(2). MLC therefore constitutes a crucial node that integrates and channels activatory and inhibitory stimuli towards myosin II function.

MLCP is composed of a 38-kDa protein phosphatase catalytic subunit (PP1c; specifically the β isoform), a 130-kDa myosin-phosphatase targeting subunit 1 (MYPT1) and, in smooth muscle, a 20-kDa subunit of unclear function (3)(4). MYPT1 acts as a scaffold that brings PP1c in close proximity to its substrates. The activity, protein-protein interactions and localization of MLCP are fine-tuned by the phosphorylation status of MYPT1, which is targeted by multiple protein kinases. Although more than 30 phosphorylation sites have been identified, the best-characterized ones are S695, T696, S852 and T853 (1). Phosphorylation of these sites by the relative activities of Rho kinase (ROCK) and cAMP and cGMP-dependent protein kinases (PKA and PKG) determines MLCP activity (1). Activation of G protein coupled receptors by a variety of agents results in ROCK-dependent MYPT1 phosphorylation at T696 and T853. This decreases the phosphatase activity of MLCP by an autoinhibitory mechanism (5)(6). Phosphorylation by PKA and PKG at the S695 and S852 sites is believed to prevent or oppose the inhibition of MLCP elicited by ROCK phosphorylation and is frequently described as disinhibition (7).

In smooth muscle cells, PKG1α binds to the C-terminal leucine zipper (LZ) motif of MYPT1 and the formation of this signaling complex is thought to be critical for the ability of nitric oxide (NO) to control smooth muscle cell tone (8). A similar mechanism has been proposed to mediate the inhibitory effects of NO in platelets (9). cAMP signaling promotes endothelial cell monolayer barrier integrity, is the most potent endogenous mechanism for the control of platelet function and is also an important mediator of smooth muscle relaxation (7)(10) (11)(12). The most important effector of cAMP is PKA. This enzyme is a heterotetramer composed of two regulatory (PKA-R) and two catalytic (PKAcat) subunits. Four PKA-R variants exist (RIα, RIβ, RIIα and RIIβ) that are differentially expressed and give rise to PKA isoforms with distinct biochemical properties and subcellular localization, possibly playing non-redundant roles in the regulation of cell functions (13).

PKA isoforms are tethered to different subcellular regions and brought to the vicinity of PKA substrates by binding to A kinase anchoring proteins (AKAPs) that define distinct signaling compartments and assemble complex signalosomes (14). AKAPs typically possess an amphipathic helix that mediates the interaction with the dimerization and docking (D/D) domain at the N-terminus of PKA-R subunits (15). In a previous study we have presented evidence for cAMP signaling compartmentalization in platelets, where moesin, the first functionally validated AKAP, targets type I PKA to lipid rafts (16). Because MYPT1 is part of a complex signaling node that includes PP1c, kinases, and other enzymes involved in signal transduction (3), we hypothesized that MYPT1 might function as an AKAP to target PKA to the MLCP signaling node, where the kinase phosphorylates MYPT1 and potentially other proteins in the complex. Using a combination of biochemical approaches, we show that MYPT1 interacts with all four PKA-R variants, that the interaction does not require the D/D domain of PKA-R but requires the PKA and ROCK substrate motif of MYPT1 around S695/T696. We propose that MYPT1 functions as a non-canonical AKAP that prevents PKAcat, and potentially also ROCK, from accessing MYPT1.

## Materials and Methods

### Antibodies

Primary antibodies against following proteins were used: MYPT1 (rabbit IgG mAb clone D6C1, #8574) from Cell Signaling Technology (Leiden, The Netherlands), MYPT1 (mouse IgG1 mAb clone C-6, sc-514261) from Santa Cruz Biotechnology (Heidelberg, Germany); mouse monoclonals against PKA regulatory subunits I (610166, IgG2b clone 18), IIα (612243, IgG1 clone 40), IIβ (610626, IgG1 clone 45), and catalytic-α subunit (610980, IgG2b clone 5B) from BD biosciences (Wokingham, UK); mouse monoclonal against β-actin (clone AC-15, Ab6276) from Abcam (Cambridge, UK); mouse monoclonal against GAPDH (clone 6C5, CB-1001) from Calbiochem/Merck (Dorset, UK); horse radish peroxidase (HRP)-conjugated anti-6His antibody (HRP-66005) from Proteintech (Manchester, UK); anti Myc mouse monoclonal 9E10 and anti-GST polyclonal antibodies were kind gifts of Angelika A. Noegel, University of Cologne, Germany. The specificity of MYPT1 and PKA subunit antibodies is shown in Suppl. Fig. 1.

Rabbit IgG and mouse IgG1 isotype immunoglobulins and total rabbit immunoglobulins were from Cell Signaling Technology (Leiden, The Netherlands). Mouse IgG2b isotype immunoglobulin was from BD Biosciences (Wokingham, UK). Secondary antibodies Alexa Fluor 568-conjugated (A11031) or Alexa Fluor 488-conjugated (A11029) donkey anti-mouse or Alexa Fluor 568-conjugated (A11036) or Alexa Fluor 488-conjugated (A11034) donkey anti-rabbit immunoglobulins (Molecular Probes, ThermoFisher Scientific, Altrincham, UK) were used for immunofluorescence. Peroxidase-conjugated (Merck, Dorset, UK) or IRDye 680 and IRDye 800-conjugated (Li-Cor Biosciences, Lincoln, USA) anti-mouse and anti-rabbit immunoglobulins were used for western immunoblot.

### Plasmids

A list of all constructs used in this study is shown in Suppl. Table 1. Human MYPT1 and PKA-R and fragments thereof were amplified by PCR using Q5^®^ High-Fidelity DNA polymerase (New England Biolabs, Herts, UK) and subcloned into appropriate vectors for expression as Myc tag fusions in mammalian cells or for expression in *E. coli* as GST or His tag fusions using standard techniques. MYPT1 (IMAGE:40008469), PKA-RIβ (IMAGE:5247772) and PKA-RIIβ (IMAGE:30331962) clones were purchased from Sino Biological Inc. (Beijing, China). Plasmids containing the coding region of PKA-RIα and PKA-RIIα were provided by Kjetil Taskén, University of Oslo. All plasmids were verified by sequencing (Eurofins Genomics, Cologne, Germany).

### Other reagents

InCELLect™ AKAP St-Ht31 inhibitor peptide was from Promega (Chilworth, UK). Human fibrinogen was from Enzyme Research (Swansea, UK). Latrunculin B was from Enzo Life Sciences (Exeter, UK). All other chemicals were obtained from Merck (Dorset, UK) or Melford Laboratories (Ipswich, UK).

### Isolation of washed platelets

Human blood was taken after informed consent from healthy, drug-free volunteers under approval of the Hull York Medical School ethic committee. Platelets were prepared as described previously (17). All platelet preparations were conducted at room temperature, and platelets were incubated for at least 30 minutes at 37°C prior to experiments.

### Cell culture and transfection

293T human embryo kidney (HEK293T) cells were grown in Dulbecco’s modified Eagle’s medium (4.5 g/l glucose) (Merck, Dorset, UK) enriched with 10% fetal bovine serum (FBS), 2 mM glutamine, 1 mM sodium pyruvate, 100 U/ml penicillin, and 100 μg/ml streptomycin. Pooled HUVECs (C-12208, lot 447Z015; Promocell, Heidelberg, Germany) were cultivated in endothelial cell growth medium MV2 supplemented with 5% (v/v) FBS, 5 ng/ml epidermal growth factor, 10 ng/ml basic fibroblast growth factor, 20 ng/ml insulin-like growth factor-1, 0.5 ng/ml vascular endothelial growth factor 165, 1 µg/ml ascorbic acid and 0.2 µg/ml hydrocortisone (Promocell, Heidelberg, Germany). HEK293T cells and HUVECs were cultivated at 37°C in a humidified incubator supplied with 5% CO_2_

HEK293T cells were transiently transfected using Lipofectamine 2000 (Invitrogen, ThermoFisher Scientific, Loughborough, UK) according to the manufacturer’s instructions and cultivated for 24 h.

### Recombinant protein expression and purification

GST and His-tagged recombinant proteins were expressed and purified from *E. coli* strains XL-1 blue, M15(pREP) or Rosetta2 using standard procedures. Briefly, GST fusions were purified by incubating the bacterial soluble fraction with pre-equilibrated glutathione Sepharose bead slurry (GE Healthcare, Munich, Germany) for 1 hour at 4°C and eluting with 10 mM reduced glutathione. His-tag fusions were purified by incubating the bacterial soluble fraction with pre-equilibrated Ni-NTA agarose bead slurry (Qiagen, Hilden, Germany) for 1 hour at 4°C and eluting with 500 mM imidazole. Purified proteins were dialyzed overnight in 20 mM Tris-HCl pH 8, 100 mM NaCl.

### GST affinity pulldown

Human washed platelets (1×10^9^/ml) were lysed with one volume of ice-cold lysis buffer (10 mM Tris-HCl pH 7.4, 150 mM NaCl, 1 mM EDTA, 1 mM EGTA, 2% Triton^®^ X-100) containing protease inhibitors and incubated for 30 min on ice. HEK293T cells transiently transfected with constructs allowing expression of Myc-tagged MYPT1 fragments were cultivated for 24 hrs at 37°C and lysed in 50 mM Tris–HCl, pH 8.0, 150 mM NaCl, 1% Triton® X-100 and protease inhibitors for 30min. Cell debris was removed by centrifugation at 10,000×g at 4°C for 10 min

PKA-R subunits and a C-terminal fragment of MYPT1were expressed as GST fusions in *E. coli* and purified by incubating the soluble fraction with 25 µl of pre-equilibrated glutathione Sepharose beads slurry for 1 hour at 4°C with rolling. After washing, 500 µg of platelet or HEK293T cell lysate, or soluble fraction of bacteria expressing His-tagged proteins were added to the beads for 2 hours at 4°C with rolling. The beads-protein complexes were pelleted by centrifugation at 1000×g for 1 min, washed with lysis buffer followed by TBST (20 mM Tris pH 7.6, 150 mM NaCl, 0.1% Tween 20) and boiled in Laemmli buffer for 5 min at 95°C. Samples were resolved by 10% SDS-PAGE. Gels were either stained with Coomassie brilliant blue or immunoblotted for the proteins of interest using standard procedures.

To calculate the amount of MYPT1 pulled down by PKA-R subunits fused to GST, the density of each MYPT1 band was divided by the density of the MYPT1 band in the lysate and by the intensity of the corresponding GST fusion in the Coomassie-stained gel, and expressed as arbitrary units.

### Immunoprecipitation and cAMP affinity pulldown

HUVECs and washed human platelets were lysed with ice cold immunoprecipitation lysis buffer (10 mM Tris-HCl pH 7.4, 150 mM NaCl, 5 mM EDTA, 10% glycerol, 1% NP-40) containing protease and phosphatase inhibitors (cOmplete™ and PhosTOP™, Roche/Merck, Dorset, UK), 1 mM sodium orthovanadate and 1 mM phenylmethylsulfonyl fluoride and incubated for 30 min on ice.

Concentrations of antibodies used for immunoprecipitation were tested by protein determination and SDS-PAGE (Supplemental Fig. 2A). Lysates (500 µg of protein) were incubated with 2 µg of anti-MYPT1 rabbit antibodies or isotype specific IgG for 1 hour at 4°C with rolling. 10 µl of pre-equilibrated protein A-coated Dynabeads (ThermoFisher Scientific, Loughborough, UK) were incubated with the lysate-antibody complex overnight at 4°C with rolling. The beads-antibody-protein complexes were sedimented with a magnetic separator rack, washed 3 times with RIPA buffer and boiled in Laemmli buffer for 3 min at 100°C. Immunocomplexes were resolved by 10% SDS-PAGE and immunoblotted using standard procedures.

Immunoblots were imaged using a ChemiDoc XRS+ Imaging System (Bio-Rad Laboratories, Watford, UK) or a LI-COR Odyssey CLx Imaging System (LI-COR Biosciences, Lincoln, USA). Images were acquired at several exposure times within the linear range of the signal. Protein band densities were analyzed with Image Lab software (Bio-Rad Laboratories, Watford, UK).

To calculate immunoprecipitation depletion ratios, first the density of each β-actin normalized band in the post-beads lysates were calculated. Next, the normalized density in the lysate incubated with anti-MYPT1 antibody was divided by the normalized density in the lysate incubated with IgG. The result was subtracted from 1 to produce the depletion ratio. To calculate co-immunoprecipitation ratios, the band intensities of the immunoprecipitation lanes were normalized to their respective IgG light chain signal. Next, the normalized values for the IgG and anti-MYPT1 lanes were added and the value of each lane was divided by the sum.

For cAMP affinity pulldown, lysates (500 µg of protein) were incubated with 25 µl of pre-equilibrated 8-(6-aminohexylamino)adenosine-3’,5’-cyclic monophosphate (8-AHA-cAMP) agarose beads or control ethanolamine agarose beads (Biolog, Bremen, Germany) overnight at 4°C with rolling. Next day beads-protein complexes were pelleted by centrifugation at 1000×g for 1 min, washed with lysis buffer followed by TBST and boiled in Laemmli buffer for 5 min at 95°C. Samples were resolved by 10% SDS-PAGE and immunoblotted for the proteins of interest using standard procedures.

The depletion ratios for the cAMP affinity pulldown were calculated as for the immunoprecipitation. Pulldowns were calculated as the band intensities of the eluate lanes divided by the β-actin normalized intensity of the respective protein in the total lysate.

### Immunoflourescence

HUVEC were processed for immunofluorescence using established protocols (18). HUVECs were seeded at 5,000-10,000 cells per well in 8-well Lab-Tek^®^ chamber slides (Nunc^®^, Merck, Dorset, UK) and grown to 60-70% confluence, fixed with 4% paraformaldehyde (PFA), permeabilized and blocked with 0.1% Triton^®^ X-100, 10% donkey serum in PBS for 30 min. Cells were immunostained overnight at 4°C with the indicated primary antibodies or isotype specific immunoglobulin (Suppl Fig. 2A, B) followed by the corresponding secondary antibodies for 45 min diluted in PBS containing 0.1% Triton^®^ X-100 and 2% donkey serum. Coverslips were mounted onto the wells using Vectashield® antifade mounting medium with DAPI (2BScientific, Kidlington, UK).

Washed platelets in suspension were fixed with an equal volume of ice-cold 4% PFA in PBS for 7 min and allowed to sediment for 1 h on poly-L-lysine (0.01% in PBS) coated coverslips. For spread platelets, coverslips were coated overnight at 4°C with 100 μg/ml fibrinogen and blocked with 5 mg/ml heat-denatured fatty-acid-free bovine serum albumin (BSA) for 1 h. Washed platelets were allowed to spread for 1 h at 37°C, then fixed with 4% PFA. Fixed platelets were permeabilized with 0.3% Triton^®^ X-100 in PBS for 5 min, stained for 1 h at room temperature with the indicated primary antibodies followed by the corresponding secondary antibodies diluted in PBG (0.5% BSA, 0.05% fish gelatine in PBS) and mounted on glass slides with gelvatol (Merck, Dorset, UK).

Slides were imaged with a ZEISS LSM710 confocal microscope equipped with a 20×/0.5 EC-Plan-NEOFLUAR objective or a ZEISS ApoTome.2 fluorescence microscope equipped with an AxioCam 506 and a Plan-Apochromat 63×/1.4 oil immersion objective (ZEISS, Cambridge, UK). Images were processed with ZEISS Zen software.

For co-localization analysis, regions of interest (125×125 pixel in the case of HUVECs; individual cells in the case of platelets) were extracted from double staining images and were used to calculate the Pearson’s correlation coefficient with the ImageJ plugin JACoP (19). A set of images in which one of the channels was rotated 90° was used as a negative control.

### In situ proximity ligation assay

PLAs were carried out using a Duolink^®^ *in situ* red starter kit mouse/rabbit (Merck, Dorset, UK). HUVECs and spread platelets were fixed, permeabilized and blocked as described above for immunofluorescence. Cells were then incubated with matching concentrations of the relevant primary antibody or isotype specific immunoglobulin (mouse and rabbit) pairs (Supplemental Fig.2A, B). This was followed by incubation with anti-mouse and anti-rabbit probes and fluorescence-producing reaction as per the manufacturer’s instructions. The samples were incubated with fluorescein isothiocyanate (FITC) labeled phalloidin for 30 min prior to mounting with Vectashield® antifade mounting medium containing DAPI (2BScientific, Kidlington, UK). Samples were imaged as described in the previous section. For platelets, the number of dots per cell were scored from images. For HUVECs, fluorescence density was calculated using ImageJ because the high density of dots made counting impractical. For each individual HUVEC the background subtracted intensity was normalized to the cell area and the result expressed as arbitrary units.

### Peptide mapping

Peptide array experiments were performed as described previously (20). Briefly, peptides covering residues 501-706 of MYPT1were generated via automatic SPOT synthesis (21)(22). Peptides were synthesized on continuous cellulose membrane supports using 9-fluorenylmethyloxycarbonyl chemistry (Fmoc) by a MultiPep RSi Robot (Intavis, Cologne, Germany). Arrays were pre-activated in absolute ethanol followed by blocking in 5% dehydrated milk in TBST for 4 h at room temperature. MYPT1 arrays were then overlaid overnight at 4°C with 10 µg/ml of His-tagged PKA-RIβ or PKA-RIIβ diluted in 150 mM NaCl, 5% glycerol, 50 mM Tris pH 7.4. PKA-R binding to MYPT1 peptides was determined by incubating the array for 4 hours at 4°C with HRP-conjugated anti-6His antibody in TBST followed by enhanced chemiluminescence detection.

### Statistical analysis

Experimental data were analyzed with GraphPad Prism v6.0 (La Jolla, CA, USA). Data are presented as means ± standard error of the mean (SEM) of at least three independent experiments. Normality was assessed by the Shapiro-Wilk test for all experiments. Differences between groups were assessed using the appropriate parametric (Student’s t test or ANOVA) or non-parametric (Mann-Whitney U or Kruskall-Wallis) test for single or multiple comparisons as indicated in the figure legends and statistical significance taken at p ≤ 0.05.

## Results

### MYPT1 forms complexes with the PKA holoenzyme

To explore the hypothesis that MYPT1 functions as a PKA anchor by interacting with PKA-R subunits, we conducted an affinity pulldown assay on platelet lysates using GST-tagged PKA-R subunits (Fig. 1A). All four PKA-R variants were able to significantly pull down MYPT1 and, as expected, also the PKAcat subunit (Fig. 1A), indicating that MYPT1 is able to form complexes with the PKA holoenzyme. We then aimed at confirming the existence of such complexes using an immunoprecipitation approach in HUVECs and platelets. In HUVEC lysates a MYPT1-specific antibody resulted in 95% immunodepletion of MYPT1 along with the identification of statistically significant amounts of PKA-RIIα, PKA-RIIβ and PKAcat, but not PKA-RI, in the immunocomplexes (Fig. 1B). Although the same antibody was able to immunodeplete MYPT1 from platelet lysates, a conclusion about co-immunoprecipitation of PKA subunits could not be reached due to all of them being pulled down by the control immunoglobulin under the same experimental conditions used in HUVEC lysates (Suppl. Fig. 3A).

**Figure 1.**
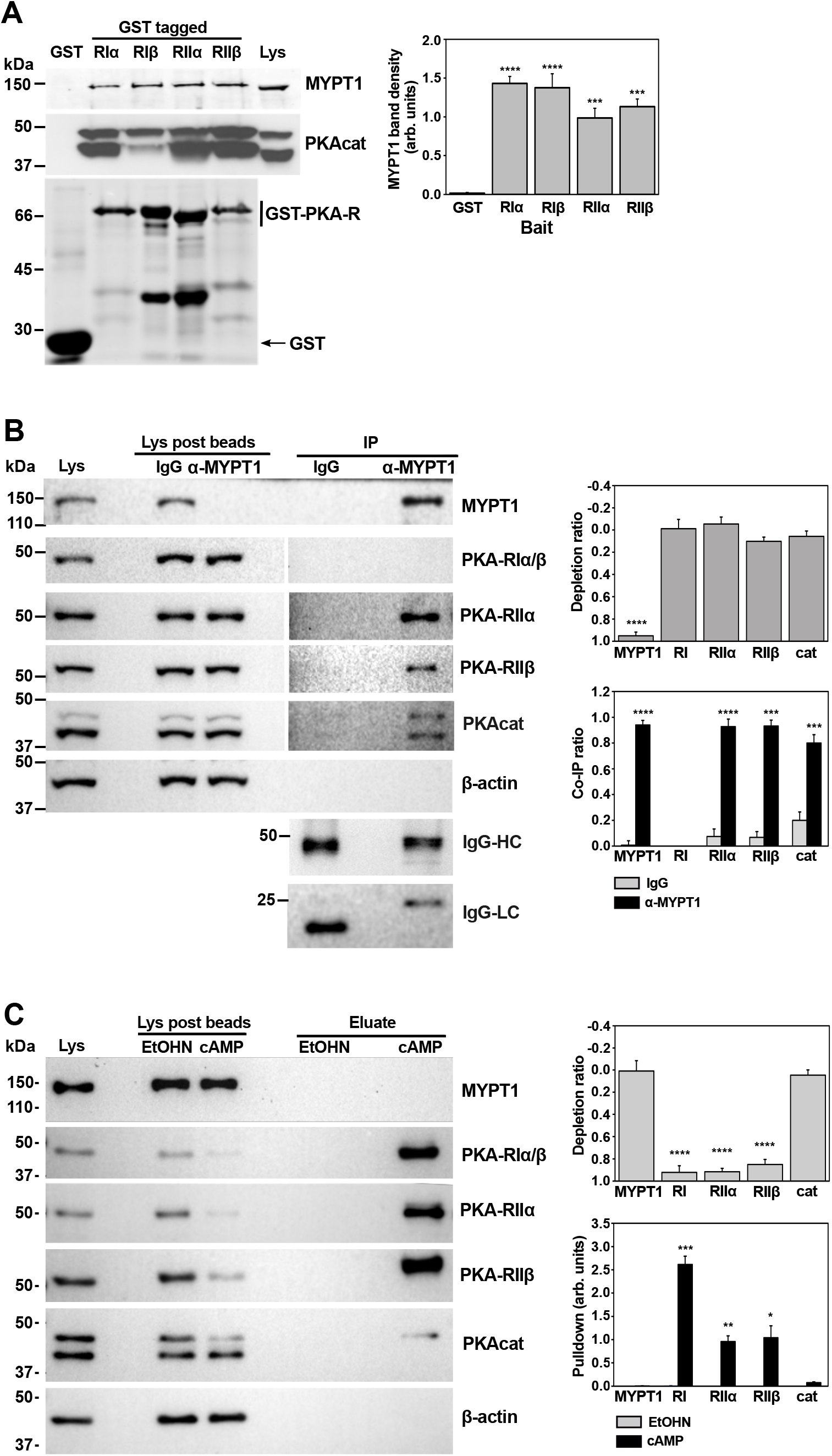
Interaction of MYPT1 with PKA in platelets and HUVECs. **(A)** Affinity pull-down with GST-tagged PKA-R subunits in platelets. Lysates were incubated with glutathione-Sepharose beads loaded with recombinant GST fusions of PKA regulatory subunits used as bait. Attached complexes were examined by western immunoblot for the presence of MYPT1 as well as for the PKA catalytic subunit. Lys, total cell lysate. The bottom panel is a Coomassie brilliant blue stained gel to analyze GST and the GST fusion proteins attached to the beads. The amounts of MYPT1 pulled down were calculated as described in Materials and Methods. (**B)**. Immunoprecipitation with anti-MYPT1 antibody in HUVECs. Lysates were subject to immunoprecipitation with rabbit MYPT1-specific antibodies or immunoglobulin G (IgG) of the same isotype. Protein complexes were examined by western immunoblot for the presence of PKA regulatory and catalytic subunits and for the IgG heavy chain (HC) and light chain (LC). Beta-actin (β-actin) was used as a loading control for lysates. Lys post beads are the lysates after incubation with and removal of the antibodies. Note that the immunoprecipitation (IP) blots for PKA subunits required longer exposures than the lysates. Depletion and co-immunoprecipitation (Co-IP) ratios were calculated as described in Materials and Methods. **(C)** Affinity pull-down with cAMP beads in HUVECs. Lysates were incubated with 8-AHA-cAMP (cAMP) or control ethanolamine (EtOHN) agarose beads. Complexes were examined by western immunoblot for the presence of MYPT1 and PKA subunits. Beta-actin (β-actin) was used as a loading control for lysates. Depletion ratios and pulldown amounts were calculated as described in Materials and Methods. Data are mean ± SEM of 3-6 independent experiments. *** p<0.001; **** p<0.0001, ANOVA followed by Tukey test (A and depletion ratios of B and C) or Student’s t test (co-IP and pulldown ratios of B and C).

Upon binding to cAMP, the PKA-R subunits change conformation and release the PKAcat subunits (13). To investigate whether the interaction of PKA with MYPT1 is conformation-dependent we performed an affinity pulldown in HUVEC and platelet lysates with agarose beads functionalized with cAMP (Fig. 1C). In HUVECs this approach depleted >85% of all PKA-R variants and, as expected, very little PKAcat was found attached to the beads, where PKA-R subunits were significantly recovered. No MYPT1 was recovered with the beads, suggesting that the PKA-MYPT1 complex occurs when PKA-R subunits are not bound to cAMP (Fig. 1C). Notably, a cAMP affinity pulldown in platelet lysates yielded a significant amount of MYPT1 bound to the beads, but also of β-actin (Suppl. Fig. 3B), in contrast to HUVEC lysates. To rule out that the interaction of MYPT1 with PKA in platelets is mediated by the actin cytoskeleton, we used latrunculin B to disrupt the actin cytoskeleton prior to the cAMP pulldown. However, under those conditions we still recovered significant amounts of MYPT1 and actin with the beads (Suppl. Fig. 3C). The different behavior of MYPT1 between HUVECs and platelets in immunoprecipitation and cAMP pulldown assays might be explained by differences in their relative amounts of actin, which are considerably higher in platelets (Suppl. Fig. 1A). MYPT1 is known to interact with actin and numerous actin-binding proteins (3) and at least PKA-RIIα has been shown to interact directly with actin too (23).

### MYPT1 co-localizes with PKA-R subunits at the cell cortex

Having shown that MYPT1 forms complexes with PKA-R subunits, we investigated whether they the colocalized in HUVECs and platelets. HUVECs were fixed and double immunostained with antibodies against MYPT1 and either PKA-I, PKA-RIIα or PKA-RIIβ (Fig. 2A). MYPT1 displayed a diffuse distribution with accumulation in the nucleus and enrichment in discrete cortical regions (Fig. 2A), as reported by others (24). The anti-PKA-RI antibody yielded a predominantly cytoplasmic pattern with very little signal in the nucleus and, in some cells, enrichment in perinuclear and cortical regions. PKA-RIIα staining was diffuse across the cytoplasm and nucleus, with a weak accumulation in cortical regions. PKA-RIIβ was also diffuse across the cell, with a strong accumulation in a paranuclear region that resembles the Golgi and, in some cells, cortical enrichment. Due to the predominantly diffuse distribution of these proteins and the variety of patterns of subcellular distribution, a noticeable colocalization of MYPT1 and PKA-R subunits was only apparent in restricted regions, predominantly at the cell cortex (Fig. 2A). Correlation analysis showed Pearson’s correlation coefficients significantly above the control (0.018 ± 0.020) for PKA-RI (0.667 ± 0.011), PKA-RIIα (0.833 ± 0.012) and PKA-RIIβ (0.640 ± 0.017) (p<0.0001) (Fig. 2B). The Pearson’s correlation coefficient for PKA-RIIα was significantly higher than that of PKA-RI and PKA-RIIβ (p<0.0001), likely as a result of the more uniform distribution of this subunit.

**Figure 2.**
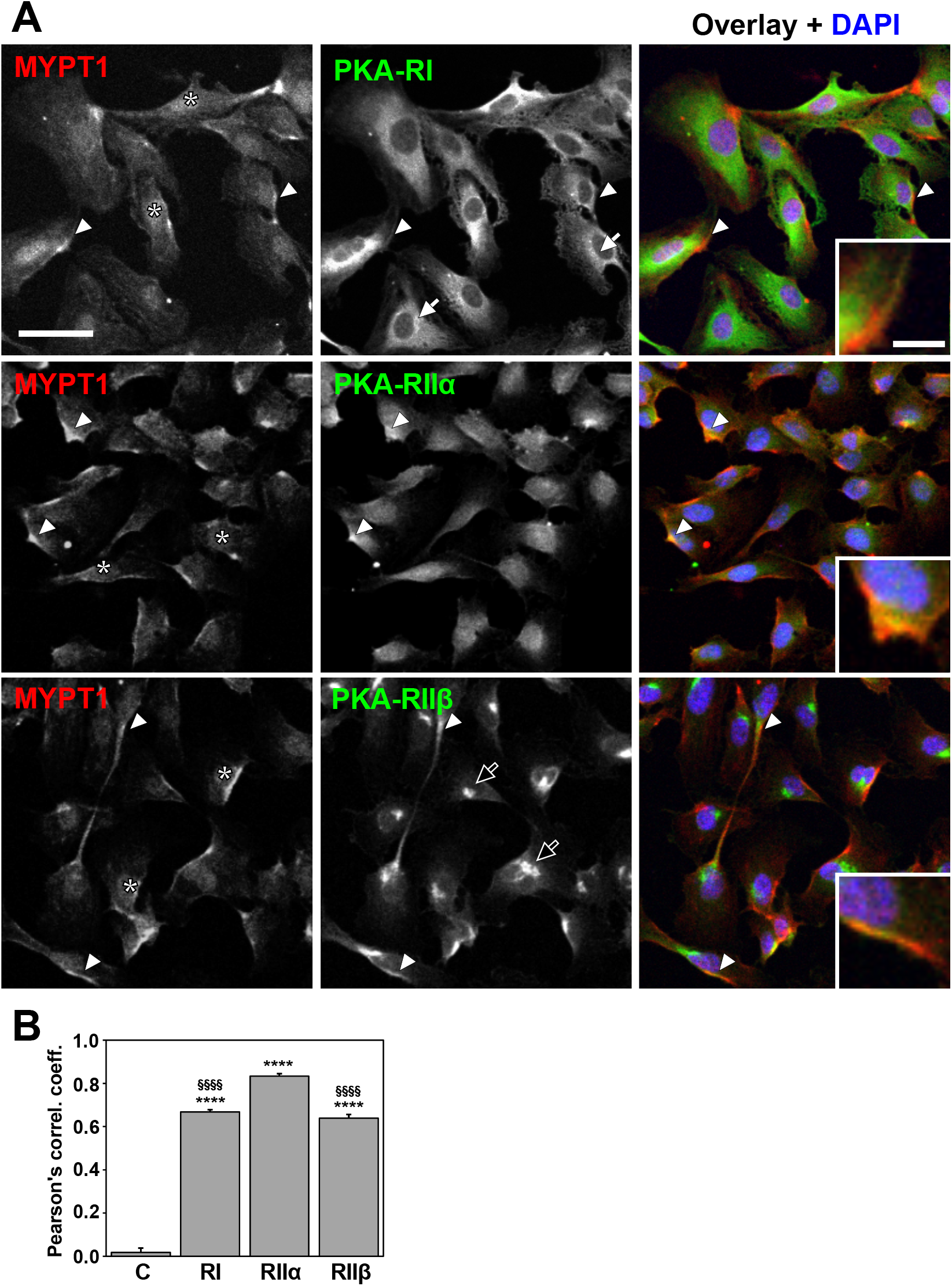
Double immunofluorescence staining of MYPT1 and PKA-R subunits in HUVECs. **(A)** Cells were fixed, permeabilized and incubated with rabbit anti-MYPT1 antibody and the indicated mouse PKA-R subunit antibody followed by species-specific fluorescently labeled Alexa fluor568 (red) or Alexa fluor488 (green) conjugated secondary antibodies. Nuclei were counterstained with DAPI. White arrowheads indicate instances of co-localization; for each image, one is zoomed in the insets. Asterisks indicate nuclear accumulation of MYPT1. White arrows point at perinuclear staining of PKA-RI. Black arrows point at Golgi-like staining of PKAR-II β. Images were acquired with a ZEISS LSM 710 confocal microscope equipped with a 20× objective. Representative images are central slices from Z stacks taken at 1.80 µm intervals. Scale bars represent 50 µm (large images) or 10 µm (insets) and apply to all images of the same size. **(B)** Pearson’s correlation coefficients for MYPT1 and PKA-R subunit double stainings. For each condition data are mean ± SEM of 30 regions of interest from 3 independent images. Images were analyzed as described in the Materials and Methods section. **** p<0.0001 relative to the control (C); §§§§ p<0.0001 relative to PKA-RIIα (Kruskal-Wallis test followed by Dunn’s test).

Instances of co-localization of MYPT1 with PKA-R subunits were also apparent in platelets (Suppl. Fig. 4A). In these cells, MYPT1 presented a diffuse punctate distribution. PKA-RI was predominantly cortical, whereas the pattern of PKA-RII variants was punctate across the cell, PKA-RIIβ being less conspicuously punctate than PKA-RIIα (16). Correlation analysis showed Pearson’s correlation coefficients significantly above the control for all PKA-R subunits in platelets (Suppl. Fig. 4B).

### PKA interacts with a central region of MYPT1 spanning amino acids 501-706

We next mapped the region of MYPT1 responsible for the interaction with PKA-R using a pull-down approach on HEK293T cells. Full length (FL) MYPT1 was divided into two major fragments: N-terminal (NT, amino acids 1-330) and C-terminal (CT, amino acids 327-1030) (Fig. 3A). Lysates of HEK293T expressing Myc-tagged FL, NT and CT MYPT1 were subjected to pull-down with GST-tagged PKA-R variants bound to glutathione Sepharose beads. FL and CT-MYPT1 co-purified with all GST-PKA-R variants (Fig. 3B). In order to further narrow down the binding region of MYPT1 with PKA-R, the CT fragment was subdivided into four overlapping fragments designated as C1, C2, C3 and C4. Each fragment was 200 amino acids long with 30 overlapping residues (Fig. 3A). Lysates of HEK293T cells expressing these Myc-tagged fragments were subjected to pull-down assays with GST-PKA-R variants as above. Only the C2 fragment was strongly associated with PKA-R variants (Fig. 3C). The C2 fragment contains the central insert (CI) region of MYPT1 as well as the S695 and T696 residues. Weak binding was observed with the C1 fragment, suggesting that a binding motif may reside in the C2 region that overlaps with C1.

**Figure 3.**
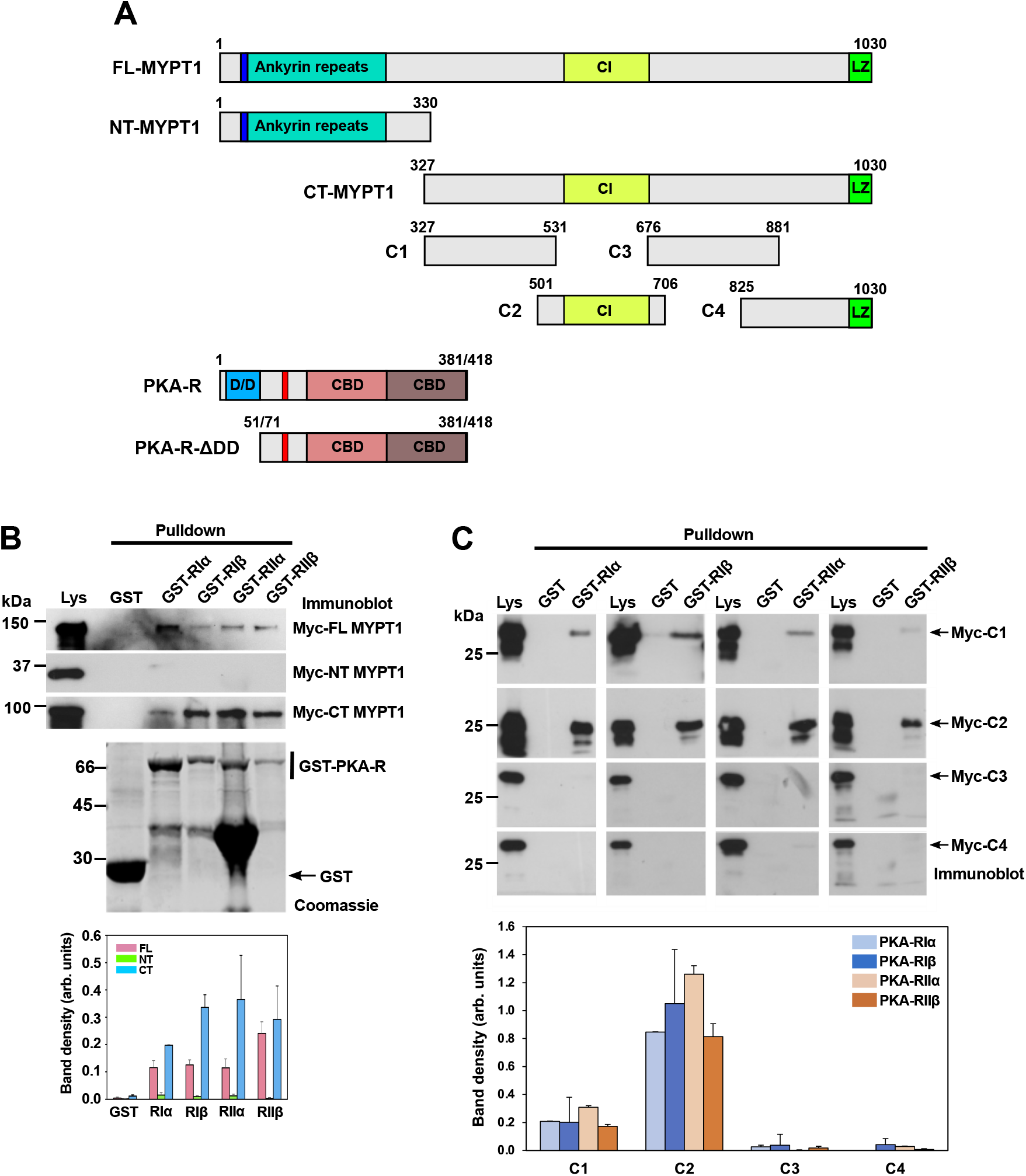
Mapping the interaction region of MYPT1 with PKA-R. **(A)** Domain architecture of the proteins and fragments used in this study. MYPT1 is composed of an N-terminal domain with PP1c binding site (RVXF, in dark blue), followed by 8 ankyrin repeats, a coiled-coil C-terminal domain containing a central insert (CI) region and a leucine zipper (LZ) motif. FL, full length protein, NT, N-terminal fragment, CT, C-terminal fragment. All PKA regulatory subunits share a domain architecture, although their total length is variable: N-terminal dimerization and docking domain (D/D), followed by the inhibitor site (in red) and two cyclic nucleotide binding domains (CBD). ΔDD are PKA-R fragments that lack the D/D domain. Different tags (Myc, GST, His) were attached to the N-terminus of the recombinant proteins (see Supplementary Table 1). **(B)** HEK293T cells were transiently transfected with constructs (FL, CT and NT) allowing expression of Myc-tagged MYPT1. After 24 hours cell lysates were incubated with glutathione-Sepharose beads previously loaded with recombinant GST fusions of PKA-R subunits. Attached complexes were examined for the presence of the indicated Myc-tagged MYPT1 fragments by immunoblot using anti-Myc antibody. Lys, lysate. Immunoblots are representative of two independent experiments. One gel was stained with Coomassie brilliant blue to analyze the GST fusion proteins attached to the beads. Protein band densities were normalized to the corresponding lysate and to the amount of GST fusion. **(C)** HEK293T cells were transiently transfected with constructs allowing expression of Myc-tagged MYPT1 C-terminal fragments. Lysates were processed and analyzed as in B. Immunoblots are representative of two independent experiments. Data in the bar diagrams of B and C are mean ± SEM.

### MYPT1 interacts directly with PKA-R isoforms

To verify that the interaction between MYPT1 and PKA-R subunits is direct, we used an affinity pulldown approach with recombinantly expressed proteins. The GST-tagged MYPT-C2 fragment was immobilized on glutathione Sepharose beads and incubated with bacterial supernatants containing His-tagged PKA-RIβ and RIIβ as representatives of type I and type II regulatory subunits. Both PKA-R variants were pulled down by the GST-MYPT1-C2 fragment, whereas no association was observed with GST (Fig. 4A), suggesting that MYPT1 and PKA-R interact directly and MYPT1 might be functioning as an AKAP. AKAPs characteristically possess an amphipathic helix that mediates the interaction with the D/D domain of PKA-R (15). We analyzed the MYPT1 amino acid sequence of MYPT1 for the presence of a possible amphipathic helix using HeliQuest (25). The analysis identified two regions, amino acids 268-285 and amino acids 987-1004, with a strong amphipathic character comparable to known AKAP disruptor peptides like Ht31 and RIAD (Suppl. Fig. 5). These two regions localize, respectively, in the N-terminal and C4 fragments tested in the previous section, none of which appears to interact with PKA-R variants.

**Figure 4.**
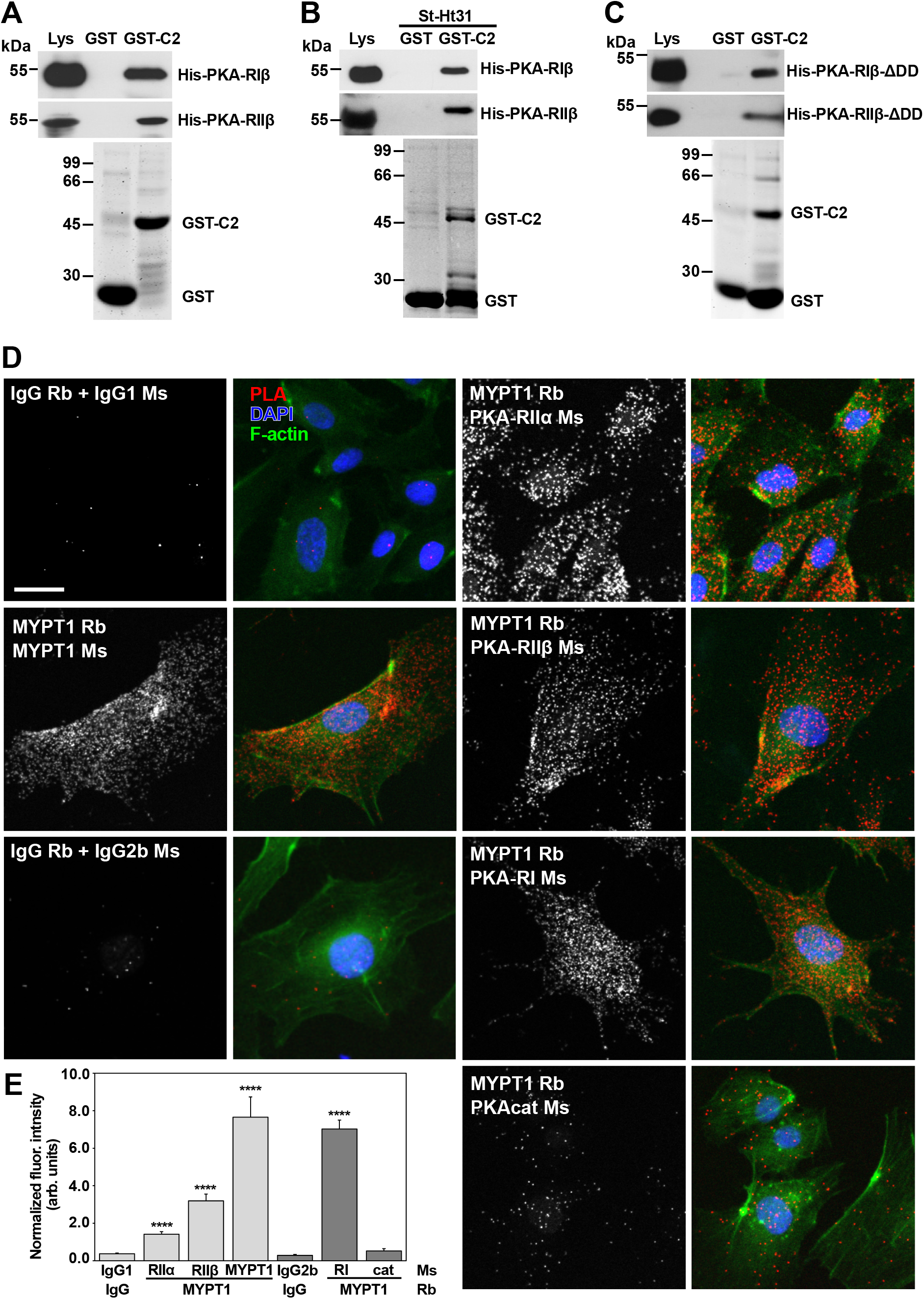
Investigating direct interaction of MYPT1 and PKA-R. **(A)** Direct interaction of MYPT1 with PKA-R subunits was tested using an affinity pulldown approach. Bacterial supernatants containing recombinant His-tagged PKA-RIβ or RIIβ were incubated with glutathione Sepharose beads previously loaded with GST or GST-tagged MYPT1-C2 (GST-C2). Ly, bacterial lysate. Protein complexes were examined by western immunoblot using anti-His antibody. The bottom gel was stained with Coomassie brilliant blue to confirm the presence of GST fusion proteins attached to the beads. **(B)** Effect of an AKAP disruptor peptide on the interaction of MYPT1 with PKA-R. The experiment was conducted as in A, with the difference that the bacterial supernatants were supplemented with 2 µM Ht31for 20 min prior to addition of the beads. **(C)** D/D domain requirement for the interaction of MYPT1 with PKA-R. The experiment was conducted as in A, with the difference that bacterial supernatants containing recombinant His-tagged PKA-RIβ or RIIβ lacking the D/D domain (see Fig. 3A) were used. Immunoblots in A, B and C are representative of three independent experiments. **(D)** Proximity ligation assay (PLA) for interaction of MYPT1 with PKA subunits in HUVECs. Cells were fixed, permeabilized and incubated with the indicated antibody pairs, followed by PLA reaction (white dots in grey scale and red dots in overlay panels). The left-hand side panels are a positive control (mouse and rabbit monoclonal antibodies anti-MYPT1) and isotype-specific immunoglobulin negative controls (mouse IgG2b for anti-PKA-RI and anti-PKAcat, mouse IgG1 for anti-PKA-RIIα, anti-PKA-RIIβ and anti-MYPT1, rabbit IgG for anti-MYPT1). Matching antibody concentrations were used. F-actin was stained with FITC-phalloidin (green) and nuclei with DAPI (blue). Images were acquired with a ZEISS LSM 710 equipped with a 20× objective. Representative images are central slices from Z stacks taken at 1.80 µm intervals comprising the complete cell thickness. Scale bar represents 25 µm and applies to all panels. **(E)** Quantification of the *in situ* PLA. Data are mean ± SEM of 14-39 cells from at least 3 independent images. Images were analyzed as described in the Materials and Methods section. Results from images where mouse antibodies of the IgG1 or IgG2b isotype were used are shown in light or dark gray, respectively. **** p<0.0001 relative to the respective isotype control (Kruskal-Wallis test followed by Dunn’s test).

AKAPs interact with the D/D domain of PKA-R subunits and tether PKA to distinct subcellular locations, spatially and temporally restricting the activity of PKA (15)(26). We therefore sought to investigate whether the interaction between MYPT1 and PKA-R involves the D/D domain of PKA-R. To this end we adopted two different pull-down approaches, either using a synthetic AKAP disruptor peptide (Ht31) or using deletion mutants of the D/D domain (ΔDD). Ht31 is a non-specific AKAP disruptor peptide, which is able to uncouple the anchoring of both PKA type I and type II to AKAPs (27). To test the effect of Ht31, the bacterial supernatants containing His-tagged PKA RIβ and RIIβ were pretreated with 2 µM of the disruptor peptide prior to the pulldown with GST-tagged MYPT-C2. This concentration of inhibitor was selected based on previous studies (16). Ht31 did not disrupt the interaction of MYPT1-C2 with any of the two PKA-R subunits, suggesting that MYPT1 does not interact with PKA in a canonical AKAP manner (Fig. 4B). This was further confirmed by using His-tagged versions of PKA RIβ (71-381) and RIIβ (51-418) lacking the D/D domain in a pulldown assay with GST-tagged MYPT1-C2. MYPT1-C2 was able to pull down both, suggesting that the D/D domain is not required for the interaction with MYPT1 (Fig. 4C). All together our data shows that MYPT1 interacts directly with PKA-R but it does not follow the mechanism of interaction characteristic of canonical AKAPs.

To confirm the interaction of MYPT1 and PKA-R *in situ*, we utilized a PLA (28) in HUVECs. To ensure that the technique works specifically, positive and negative control reactions were tested. A positive control with a combination of rabbit and mouse anti-MYPT1 antibodies produced numerous dots, whereas negative controls with isotype-specific mouse and rabbit immunoglobulins showed very little signal (Fig. 4D, left panels and Fig. 4E). PLA reactions using an anti-MYPT1 rabbit antibody in combination with subunit-specific anti-PKA-R mouse antibodies exhibited numerous dots significantly above the respective negative controls, whereas the reaction with an anti-PKAcat mouse antibody yielded levels not significantly above the negative control (Fig. 4D, right panels and Fig. 4E). The signal obtained with anti-PKA-RI antibody was well above that obtained with each anti-PKA-RII antibody, probably reflecting the fact that the anti-PKA-RI antibody recognizes both PKA-RI and RIβ variants. The same approach was used in platelets spread on fibrinogen with comparable results (Suppl. Fig. 6). We scored 1.48 ± 0.14 dots per platelet with anti-PKA-RI antibody, 1.01 ± 0.11 with anti-PKA-RIIα and 0.97 ± 0.10 with anti-PKA-RIIβ, all significantly above the negative controls (0.12 ± 0.04). The reaction with PKAcat antibody (0.37 ± 0.06 dots per platelet) was not significantly above its negative control. The results suggest that a significant number of PKA-R subunits exist in a complex with MYPT1 both in HUVECs and platelets.

### Peptide array mapping narrows down the MYPT1-PKA-R interaction sites

Having demonstrated that MYPT1 interacts directly with PKA-R variants and narrowed down the interaction region down to a C-terminal fragment encompassing residues 501-706 (MYPT1-C2), we sought to map the sites of this interaction using a peptide array overlay. To this end, a peptide array covering MYPT1-C2 was overlaid with His-tagged PKA-RIβ and II β as representatives of type I and type II regulatory subunits (Fig. 5A and Suppl. Fig. 7). Both PKA-R variants demonstrated a similar pattern of binding to three regions of MYPT1-C2, with stronger binding identified to peptides 13-18 and 29-37 that may represent primary binding sites, whereas peptides 1-6 may represent an accessory binding site. Peptides 1-6 localize in the portion of MYPT1-C2 that overlaps with MYPT1-C1, which explains the relatively weak binding of MYPT1-C1 fragment to PKA-R variants. Peptides 15 and 35 appeared to be the strongest binding peptides within each binding region and were selected for follow up substitution analysis. Comprehensive alanine substitutions of these two peptides showed abolished binding when K595 was replaced by alanine, suggesting that this is a major residue involved in the interaction with PKA-R (Fig. 5B and Suppl. Fig. 8). In peptide 35, E676 is part of an acidic patch and we could observe almost abolished binding when this residue was replaced by alanine as well as a trend towards reduced binding when the adjacent D675 and E677 were replaced by alanine. However, the binding was not affected when these acidic residues were replaced by lysines (Fig. 5C and Suppl. Fig. 8). Substitution analysis of residues 693-696 revealed a 50% binding reduction when R693, R694 and S695 were replaced by alanine and abolished binding when T696 was replaced. Furthermore, substitution of R693 and R694 by residues of the opposite charge abolished binding (Fig. 5C and Suppl. Fig. 8). These results indicate that residues involved in the substrate motif for Ser/Thr protein kinases are important for the interaction.

**Figure 5.**
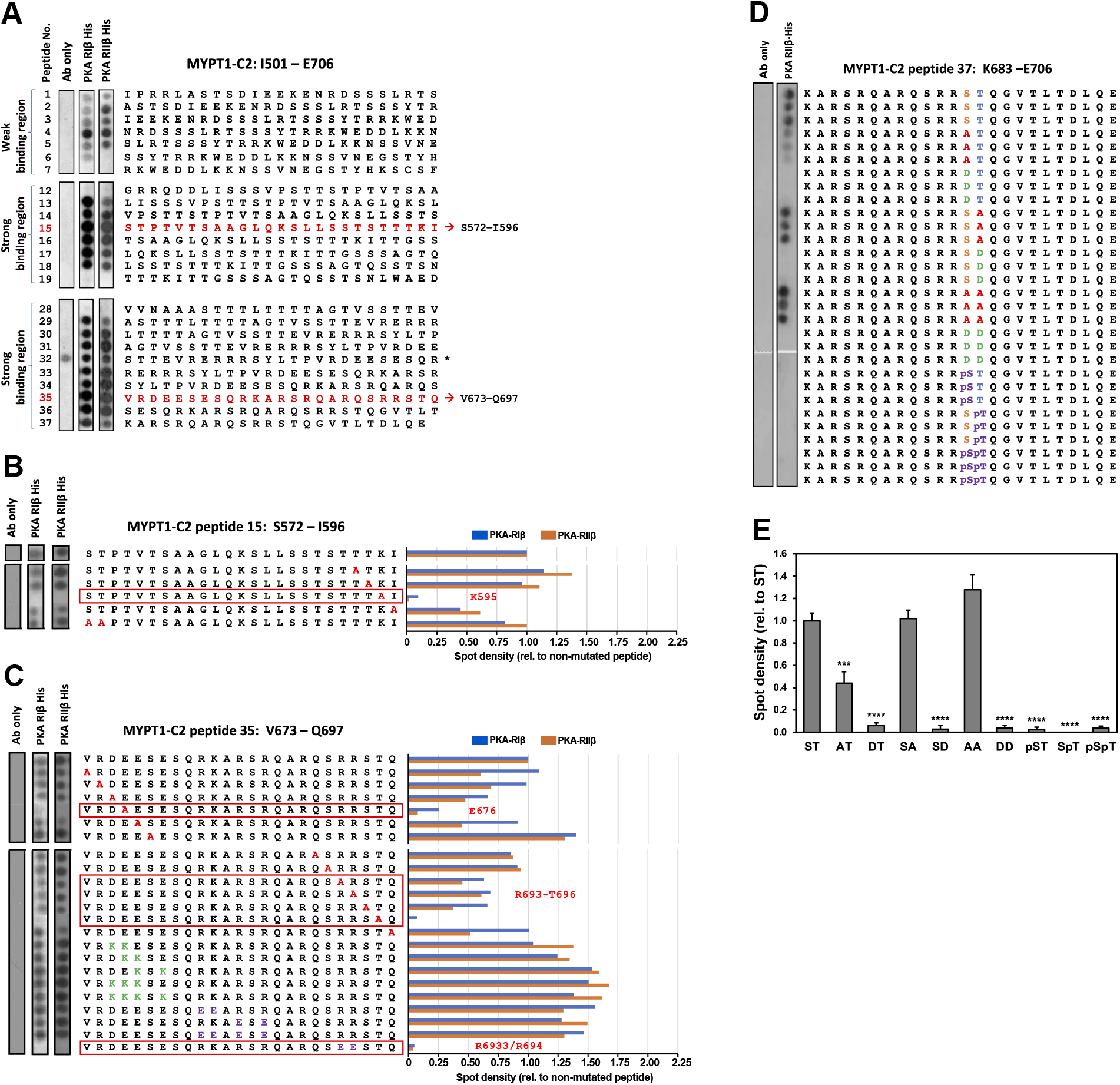
Peptide array epitope mapping of PKA-R subunits on MYPT1. **(A)** An array of overlapping 25mers, sequentially shifted by 5 amino acids, spanning MYPT1-C2 protein segment (I501 – E706) was overlaid with recombinant His-tagged PKA-RIβ or RIIβ, probed with HRP conjugated anti-polyHis antibody and visualized by enhanced chemiluminescence. For the antibody only (Ab only) control, PKA-R was omitted. Immunoblots covering the full peptide array are shown in Supplemental Fig. 6. Binding peptides selected for follow up analysis are indicated in red. The asterisk next to peptide 32 indicates antibody-associated non-specific binding. **(B)** Peptide array substitution analysis of MYPT1-C2 peptide 15. Key MYPT1-C2 hot spots were identified by single, dual or multiple alanine substitutions. Arrays were processed as in panel A. The red box highlights a peptide where a substitution resulted in complete loss of binding. Spot densities are shown relative to the non-mutated peptide after subtraction of the antibody only control. **(C)** Peptide array substitution analysis of MYPT1-C2 peptide 35. Key MYPT1-C2 hot spots were identified by single alanine substitutions or by opposite charge residue substitutions. Arrays were processed as in panel A. The red boxes highlight peptides where substitutions resulted in consistent reduction or complete loss of binding. Spot densities are shown relative to the non-mutated peptide after subtraction of the antibody only control. Immunoblots covering full arrays of peptides 15 and 35 are shown in Supplemental Fig. 7. **(D)** Peptide array substitution analysis of phosphorylation targets S695 and T696. An array of overlapping 24mers, sequentially shifted by 5 amino acids, spanning MYPT1-C2 peptide 37 was used for single and double alanine, aspartate or phosphorylated residue substitutions. The array was processed as in panel A but was probed only for PKA-RIIβ. Each substitution was tested in triplicate. **(E)** Spot densities of panel D were analyzed with Image J. For each spot the intensity of the corresponding antibody only background spot was subtracted and calculated relative to the non-mutated (ST) peptide. Data are mean ± SEM. *** p<0.001, **** p<0.0001 relative to the ST peptide, ANOVA followed by Tukey test.

We next investigated whether the phosphorylation status of S695 and T696 would affect the interaction with PKA-R. Having established that PKA-RIβ and RIIβ behave similarly, we restricted the study to PKA-RIIβ. A peptide array was synthesized in which S695 and T696 were substituted, either individually or in combination, by either alanine, the phosphomimetic aspartic acid or the corresponding phosphorylated residue. Peptide 37 was used for this purpose, to provide additional structural environment to the S695 and T696 residues (Fig. 5D, E). Under these conditions, substitution of S695 by alanine resulted in weaker binding of PKA-RIIβ, whereas substitution of T696 or both did not significantly alter binding. In contrast, substitution of S695 and T696, alone or in combination, by aspartic acid or a phosphorylated residue completely abolished binding. This indicates that the phosphorylation status of S695 and T696 might modulate the interaction of MYPT1 with PKA-R.

## Discussion

In this study we addressed the hypothesis that MYPT1 contributes to compartmentalize PKA signaling by functioning as an AKAP that would target PKA to the MLCP signaling node. This implies that MYPT1 interacts with PKA-R. Using affinity pulldowns and immunoprecipitation we show that MYPT1 is able to form complexes with the PKA holoenzyme. Furthermore, using affinity pulldowns with recombinant proteins and *in situ* PLA assays we demonstrate that MYPT1 and PKA-R subunits interact directly, supporting the conclusion that MYPT1 might function as an AKAP. Characteristically, AKAPs possess an amphipathic helix that mediates the interaction with the D/D domain of PKA-R (15); however, the two amphipathic helices we identified *in silico* in MYPT1 localize outside of the binding region uncovered in affinity pulldowns. Moreover, the interaction does not require the D/D domain of PKA-R, further ruling out MYPT1 as a canonical AKAP. While the vast majority of AKAPs recruit PKA-R subunits through a signature amphipathic helix that interacts with the D/D domain, a number of proteins display a different mode of interaction. Examples of non-canonical AKAPs are pericentrin (29), RSK1 (30), neurochondrin(31), tubulin (32), actin (23) and α4 integrin (33) and their interaction with PKA-R does not involve an amphipathic helix (34). MYPT1 fulfils this criterium to be considered a non-canonical AKAP and, additionally, its binding to PKA-R does not require the D/D domain. MYPT1 exhibits dual specificity, being capable of binding both type I and type II PKA-R subunits, although the relative affinities for each PKA-R variant remain to be established. Besides recruiting PKA holoenzymes, MYPT1 is also a PKA substrate (35). In general, AKAPs recruit PKA holoenzymes to the proximity of PKA substrates, but cases have been reported where the AKAP itself is also a PKA substrate (36). The orphan G protein coupled receptor Gpr161 is an example of a receptor that functions as high affinity AKAP for PKA-RI through an amphipathic helix at its C-terminal tail while also carrying a PKA phosphorylation site shortly upstream (36). AKAP79 is an AKAP for PKA-RII but is also a substrate for the anchored kinase (37). The main difference between these two AKAPs and MYPT1 is that in the latter the PKA phosphorylation site is part of the PKA-R binding region.

Using a peptide array overlay approach, we have narrowed the interaction of PKA-R down to three MYPT1 regions. Both PKA-RI and RII subunits appear to rely upon the same residues for effective binding to MYPT1, indicating they may compete for the same binding sites. Furthermore, substitution analysis has identified residues K595, E676 as well as the PKAcat substrate motif (R693-T696; RRST in what follows) as critical for the interaction. PEP-FOLD3 (38) predicts peptide 16, which contains K595, to be in part helical (T577-L587) (Fig. 6). Peptide 35, which contains E676 and the PKAcat substrate motif, is part of an α-helix of the S658-S714 peptide resolved by nuclear magnetic resonance (Fig. 6). Both PKA-RI and RII subunits were unable to bind to MYPT1 following dual substitution of R693 and R694 by glutamic acid, which flipped the electric charges. This suggests the existence of a direct point of contact between PKA-R and MYPT where salt bridges are established between oppositely charged residues. The phosphorylation status of S695 and T696 is also critical for the interaction of MYPT1 with PKA-R and may help explain the observation that phosphorylation of T696 by ROCK prevents the subsequent phosphorylation of S695 (35).

**Figure 6.**
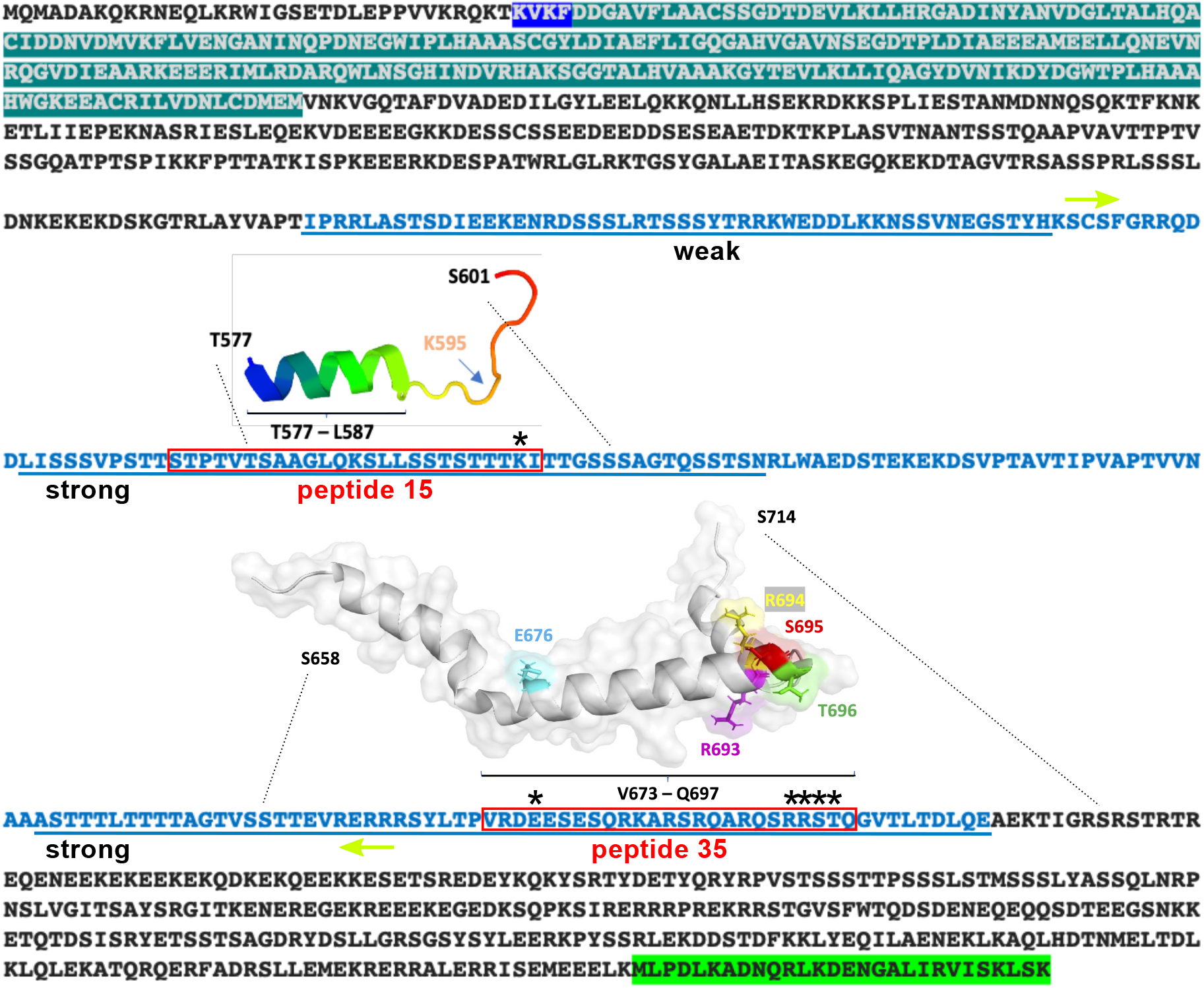
Summary of the interaction between MYPT1 and PKA-R. MYPT1 domains are highlighted: PP1c binding site (dark blue), ankyrin repeats (teal) and leucine zipper motif (green). The central insert region spans between the lime color arrows. The stretch encompassing the MYPT1-C2 region identified in GST pulldown assays (Fig. 3C) is shown in blue characters. The three binding regions identified in the peptide array overlay (Fig. 5A) are underlined. Red boxes contain peptides 15 and 35 used for substitution analysis. Asterisks indicate critical residues involved in the interaction with PKA-R identified in peptide array screening analyses (Fig. 5B-D). Structures of peptides 16 and 35 are shown. For peptide 16, PEP-FOLD3.5 predictive software indicates structure to be in part (T577-L587) helical based on 200 simulations. Note that peptide 16 was used instead of 15 in order to provide more structural context to K595. Peptide 35 is part of an amphipathic α-helix of the T658-S714 peptide resolved by nuclear magnetic resonance (PDB accession number 2KJY). Critical residues are colored in the structures.

Further studies will be necessary to define the interfaces between PKA-R and MYPT1, outside the phosphorylation sites, that determine the specificity of recognition of PKA-R as opposed to ROCK and other kinases that target the same phosphorylation site. A recent study using GST pulldowns and synthetic peptides has identified short MYPT1 regions (docking motifs) close to T696 and T853 that are important for interaction with ROCK (39). One docking motif encompasses residues 682-690 (RKARSRQAR) and is therefore part of peptide 35. While this suggests overlapping between ROCK and PKA-R interaction sites, the requirements for each kinase appear to be different: replacing basic amino acids of the 682-690 docking motif by alanine negatively affects the interaction with ROCK (39) but does not affect the interaction with PKA-R. In contrast, the acidic patch around E676 might be critical for interaction with PKA-R.

Collectively, our findings fit a model in which recognition of the RRST motif of MYPT1 by PKA happens primarily through the PKA-R subunits (Fig. 7). When the levels of cAMP increase, binding of this second messenger to PKA-R triggers a conformational change that results in dissociation of PKAcat. It is possible that the same conformational change causes the dissociation of PKA-R from MYPT1, which would explain our cAMP pulldown data where PKA-R bound to 8-AHA-cAMP functionalized beads is unable to pull down MYPT1. PKAcat would then be able to access the RRST motif and phosphorylate the S695 residue, thus shielding MYPT from ROCK. In this respect, it has been reported that prior phosphorylation of S695 by PKA *in vitro* excludes phosphorylation of T696 without affecting the activity of PP1c (7)(35). S695 becomes a substrate of the PP1c subunit of MLCP, cancelling at some point the effects of cAMP-dependent signaling and reverting MYPT1 to a state where S695 is no longer phosphorylated and it can again recruit PKA. Our PLA results suggest that, at least in HUVECs, a substantial proportion of PKA-R subunits associate to MYPT1 in the absence of any stimulus. Because experimental measurements of the cellular concentrations of PKA subunits report a 10-15-fold molar excess of PKA-R to PKAcat (40)(41), the PKA-R subunits associated to MYPT1 probably contribute to rapidly recapturing and inactivating PKAcat and maintaining a pool of catalytic subunits in the vicinity of MYPT1.

**Figure 7.**
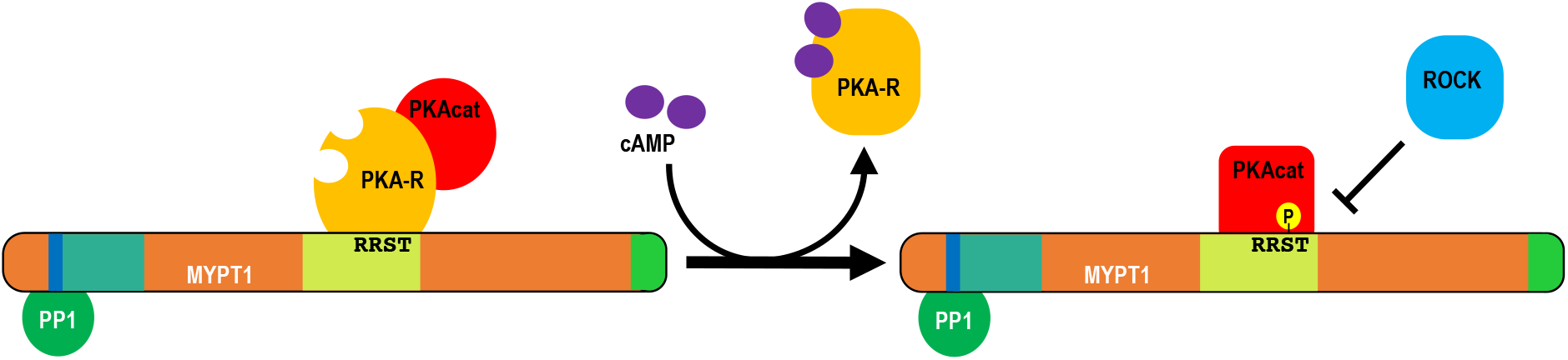
Working model of the interaction between MYPT1 and PKA-R. For simplicity, PKA subunits are shown as monomers. In resting conditions PKA-R binds to MYPT1 at the RRST motif and recruits PKAcat (PKAc), but prevents it from phosphorylating S695. Upon binding of cAMP, PKA-R changes conformation, unmasks S695 and allows phosphorylation of this residue by activated and released PKAcat. The phosphorylated RRST motif is protected from ROCK and other kinases, but PP1c activity is not affected and subsequently would dephosphorylate S695. MYPT1 would then become available for association with the PKA holoenzyme. The domain architecture of MYPT1 is indicated by colors: PP1c binding motif (blue), ankyrin repeats (teal), central insert (lime) and leucine zipper motif (green).

In summary, our findings reveal a novel mechanism of regulation of MYPT1 phosphorylation by PKA. This mechanism involves the direct interaction of PKA-R with a central MYPT1 region that includes the kinase substrate motif of both PKAcat and ROCK. We propose that MYPT1 could function as a non-canonical AKAP to target the PKA holoenzyme to the MLCP signaling node, where it would modulate the cross-talk between PKA and ROCK and MLCP activity. Future studies should address the functional relevance of the novel mechanism we have uncovered in this study.

## Acknowledgements

J.S.K. was a recipient of a University of Hull PhD scholarship and is supported by British Heart Foundation grant RG/F/22/110067. P.A.S. was a recipient of a University of Hull PhD scholarship in the cluster “Health*GDP—Health Global Data Pipeline for biomedical research and clinical applications” (2018-2023; L.L.N. – cluster lead). J.L. was supported by a British Heart Foundation PhD scholarship (PG/21/10363). W.J. was supported by the Youth Foundation of Science and Technology Department of Jilin Province (project No. 20150520033JH) and was a recipient of a Chinese Academy of Science postdoctoral scholarship.

## Notes

### Competing Interest Statement

The authors have declared no competing interest.

## References

1. J. A. MacDonald, M. P. Walsh, Regulation of smooth muscle myosin light chain phosphatase by multisite phosphorylation of the myosin targeting subunit, MYPT1. Cardiovasc. Hematol. Disord. Targets 18, 4–13 (2018).

2. M. D. Álvarez-Santos, M. Álvarez-González, S. Estrada-Soto, B. Bazán-Perkins, Regulation of myosin light-chain phosphatase activity to generate airway smooth muscle hypercontractility. Front. Physiol. 11, 1–8 (2020).

3. A. Kiss, F. Erdődi, B. Lontay, Myosin phosphatase: Unexpected functions of a long-known enzyme. Biochim. Biophys. Acta - Mol. Cell Res. 1866, 2–15 (2019).

4. R. P. Dippold, S. A. Fisher, Myosin phosphatase isoforms as determinants of smooth muscle contractile function and calcium sensitivity of force production. Microcirculation 21, 239–248 (2014).

5. A. Khromov, N. Choudhury, A. S. Stevenson, A. V. Somiyo, M. Eto, Phosphorylation-dependent autoinhibition of myosin light chain phosphatase accounts for Ca2+ sensitization force of smooth muscle contraction. J. Biol. Chem. 284, 21569–21579 (2009).

6. A. Murányi, et al., Phosphorylation of Thr695 and Thr850 on the myosin phosphatase target subunit: Inhibitory effects and occurrence in A7r5 cells. FEBS Lett. 579, 6611–6615 (2005).

7. A. A. Wooldridge, et al., Smooth muscle phosphatase is regulated in vivo by exclusion of phosphorylation of threonine 696 of MYPT1 by phosphorylation of serine 695 in response to cyclic nucleotides. J. Biol. Chem. 279, 34496–34504 (2004).

8. H. K. Surks, et al., Regulation of myosin phosphatase by a specific interaction with cGMP-dependent protein kinase lalpha. Science (80-.). 286, 1583–1587 (1999).

9. A. Aburima, K. Walladbegi, J. D. Wake, K. M. Naseem, cGMP signaling inhibits platelet shape change through regulation of the RhoA-Rho Kinase-MLC phosphatase signaling pathway. J. Thromb. Haemost. 15, 1668–1678 (2017).

10. A. Aburima, K. S. Wraith, Z. Raslan, R. Law, cAMP signaling regulates platelet myosin light chain (MLC) phosphorylation and shape change through targeting the RhoA-Rho kinase-MLC phosphatase signaling pathway. Blood 122, 3353–3345 (2013).

11. M. Aslam, et al., cAMP/PKA antagonizes thrombin-induced inactivation of endothelial myosin light chain phosphatase: role of CPI-17. Cardiovasc Res 87, 375–384 (2010).

12. R. Bátori, et al., Differential mechanisms of adenosine- and ATPγS-induced microvascular endothelial barrier strengthening. J. Cell. Physiol. 234, 5863–5879 (2019).

13. M. G. Gold, Swimming regulations for protein kinase A catalytic subunit. Biochem. Soc. Trans. 47, 1355–1366 (2019).

14. O. Torres-Quesada, J. E. Mayrhofer, E. Stefan, The many faces of compartmentalized PKA signalosomes. Cell. Signal. 37, 1–11 (2017).

15. G. Pidoux, K. Taskén, Specificity and spatial dynamics of protein kinase a signaling organized by A-kinase-anchoring proteins. J. Mol. Endocrinol. 44, 271–284 (2010).

16. Z. Raslan, S. Magwenzi, A. Aburima, K. Taskén, K. M. Naseem, Targeting of type I protein kinase A to lipid rafts is required for platelet inhibition by the 3′,5′-cyclic adenosine monophosphate-signaling pathway. J. Thromb. Haemost. 13, 1721–1734 (2015).

17. P. Joshi, et al., The membrane-associated fraction of cyclase associate protein 1 translocates to the cytosol upon platelet stimulation. Sci. Rep. 8, 10804 (2018).

18. L. L. Nikitenko, et al., Adrenomedullin and CGRP interact with endogenous calcitonin-receptor-like receptor in endothelial cells and induce its desensitisation by different mechanisms. J. Cell Sci. 119, 910–922 (2006).

19. S. Bolte, F. Cordelières, A guided tour into subcellular colocalization analysis in light microscopy. J. Microsc. 224, 213–232 (2006).

20. K. M. Brown, et al., Phosphodiesterase-8A binds to and regulates Raf-1 kinase. Proc Natl Acad Sci U S A 110, E1533–E1542 (2013).

21. R. Frank, Spot-synthesis: an easytechnique for the positionally addressable, parallel chemical synthesis in a membrane support. Tetrahedron 48, 9217–9232 (1992).

22. H. Amartely, I.-A. Anat, A. Friedler, Identifying protein-protein interaction sites using peptidearrays. J Vis Exp 18, e52097 (2014).

23. R. L. Rivard, M. Birger, K. J. Gaston, A. K. Howe, AKAP-independent localization of type-II protein kinase A to dynamic actin microspikes. Cell Motil Cytoskelet. 66, 693–709 (2009).

24. G. P. Van Nieuw Amerongen, et al., Involvement of Rho kinase in endothelial barrier maintenance. Arterioscler. Thromb. Vasc. Biol. 27, 2332–2339 (2007).

25. R. Gautier, D. Douguet, B. Antonny, G. Drin, HELIQUEST: a web server to screen sequences with specific α-helical properties. Bioinformatics 24, 2101–2102 (2008).

26. Z. Raslan, A. Aburima, K. M. Naseem, The spatiotemporal regulation of cAMP signaling in blood platelets-Old friends and new players. Front. Pharmacol. 6, Article 266 (2015).

27. A. J. Stokka, et al., Characterization of A-kinase-anchoring disrupters using a solution-based assay. Biochem. J. 400, 493–499 (2006).

28. L. U. Fredriksson S, Gullberg M, Jarvius J, Olsson C, Pietras K, Gústafsdóttir SM, Ostman A, Protein detection using proximity-dependent DNA ligation assays. Nat 20(5):.1038/nbt0502-4473-7. doi: 1073. PMID: 11981560. Nat. Biotechnol. 20, 473–477 (2002).

29. D. Diviani, L. K. Langeberg, S. J. Doxsey, J. D. Scott, Pericentrin anchors protein kinase A at the centrosome through a newly identified RII-binding domain. Curr. Biol. 10, 417–420 (2000).

30. D. Chaturvedi, H. M. Poppleton, T. Stringfield, A. Barbier, T. B. Patel, Subcellular Localization and Biological Actions of Activated RSK1 Are Determined by Its Interactions with Subunits of Cyclic AMP-Dependent Protein Kinase. Mol. Cell. Biol. 26, 4586–4600 (2006).

31. J. S. Hermann, et al., Neurochondrin is an atypical RIIα-specific A-kinase anchoring protein. Biochim. Biophys. Acta - Proteins Proteomics 1854, 1667–1675 (2015).

32. T. Kurosu, A. I. Hernández, J. Wolk, J. Liu, J. H. Schwartz, α/β-tubulin are A kinase anchor proteins for type I PKA in neurons. Brain Res. 1251, 53–64 (2009).

33. C. J. Lim, et al., α4 Integrins are Type I cAMP-dependent protein kinase-anchoring proteins. Nat. Cell Biol. 9, 415–421 (2007).

34. A. Dema, E. Perets, M. S. Schulz, V. A. Deák, E. Klussmann, Pharmacological targeting of AKAP-directed compartmentalized cAMP signalling. Cell. Signal. 27, 2474–2487 (2015).

35. M. E. Grassie, et al., Cross-talk between Rho-associated kinase and cyclic nucleotide-dependent kinase signaling pathways in the regulation of smooth muscle myosin light chain phosphatase. J. Biol. Chem. 287, 36356–36369 (2012).

36. V. A. Bachmann, et al., Gpr161 anchoring of PKA consolidates GPCR and cAMP signaling. Proc. Natl. Acad. Sci. U. S. A. 113, 7786–7791 (2016).

37. M. L.Dell’Acqua, M. C. Faux, J. Thorburn, A. Thorburn, J. D. Scott, Membrane-targeting sequences on AKAP79 bind phosphatidylinositol-4,5-bisphosphate. EMBO J. 17, 2246–2260 (1998).

38. A. Lamiable, et al., PEP-FOLD3: faster de novo structure prediction for linear peptides in solution and in complex. Nucleic Acids Res. 44, 449–454 (2016).

39. M. Amano, Y. Kanazawa, K. Kozawa, K. Kaibuchi, Identification of the Kinase-Substrate Recognition Interface between MYPT1 and Rho-Kinase. Biomolecules 12, 1–16 (2022).

40. R. Walker-Gray, F. Stengel, M. G. Gold, Mechanisms for restraining cAMP-dependent protein kinase revealed by subunit quantitation and cross-linking approaches. Proc. Natl. Acad. Sci. U. S. A. 114, 10414–10419 (2017).

41. T. T. Aye, et al., Proteome-wide protein concentrations in the human heart. Mol. Biosyst. 6, 1917–1927 (2010).

