## Supplemental tables and figures for "MYPT1 is a non-canonical AKAP that tethers PKA to the MLCP signaling node"

**Supplemental material**

**Supplemental Table 1. Plasmids used in this study.** The cloned fragments and corresponding amino acid position as well as the vector backbone are given in the table. All tags are placed at the N-terminus of the protein of interest. See also Figure 3A for a schematic of deletion constructs of MYPT1 and PKA-R.

| Insert | Amino acids | Tag | Backbone | Source |
|---|---|---|---|---|
| HsMYPT1-FL | 1-1030 | Myc | pCMV-Myc | Clontech (Takara Bio, Göteborg, Sweden). |
| HsMYPT1-NT | 1-330 | Myc | pRK5-Myc | |
| HsMYPT1-CT | 327-1030 | Myc | pCMV-Myc | |
| HsMYPT1-C1 | 327-531 | Myc | pCMV-Myc | |
| HsMYPT1-C2 | 501-706 | Myc | pCMV-Myc | |
| HsMYPT1-C3 | 676-881 | Myc | pCMV-Myc | |
| HsMYPT1-C4 | 825-1030 | Myc | pCMV-Myc | |
| HsPKARIα | 1-381 | GST | pGEX-4T-2 | Pharmacia (GE Healthcare, Munich, Germany) |
| HsPKARIβ | 1-381 | GST | pGEX-3X | |
| HsPKARIIα | 1-404 | GST | pGEX-4T-2 | |
| HsPKARIIβ | 1-418 | GST | pGEX-3X | |
| HsMYPT1-C2 | 501-706 | GST | pGEX-4T-2 | |
| HsPKARIβ | 1-381 | His | pQE32 | Qiagen (Hilden, Germany). |
| HsPKARIIβ | 1-418 | His | pQE32 | |
| HsPKARIβΔDD | 71-381 | His | pQE32 | |
| HsPKARIIβΔDD | 51-418 | His | pQE32 | |

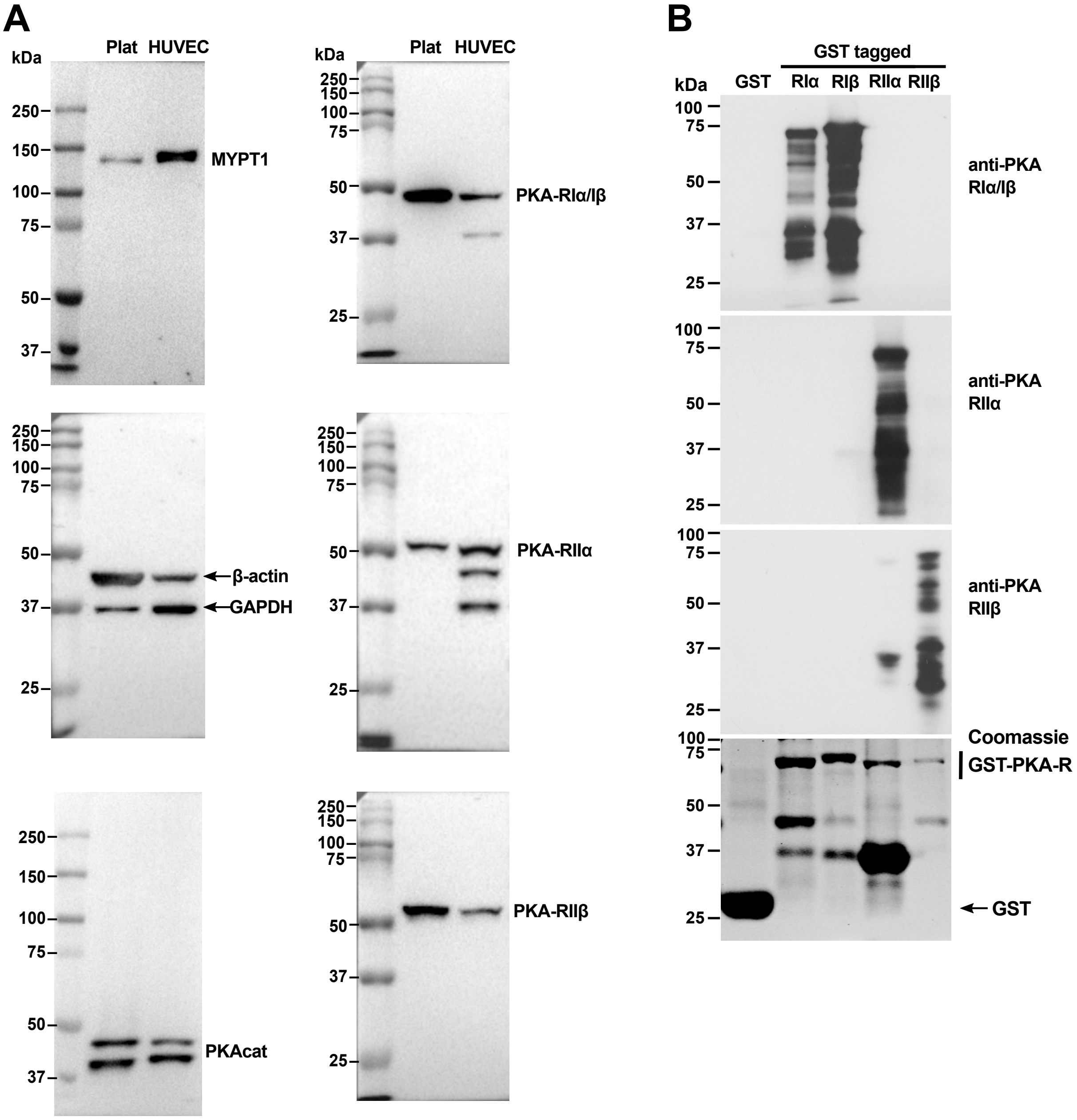


**Supplemental Figure 1. Specificity of antibodies used in this study. (A)** Human washed platelet and HUVEC lysates (40 μg) were resolved by 10% SDS-PAGE and immunoblotted with antibodies against the indicated proteins. Results with a rabbit anti-MYPT1 antibody are shown. A mouse antibody gave identical result. The anti-PKAcat antibody raised against the α variant (40 kDa) also recognizes the 93% identical slightly larger β variant. GAPDH and β-actin were used as loading controls. **(B)** PKA regulatory subunits (RIα, RIβ, RIIα and RIIβ) were expressed as GST fusions, affinity-purified on glutathione Sepharose, resolved by 10% SDS-PAGE and immunoblotted with anti-PKA RI, RIIα and RIIβ antibodies. One gel (bottom) was stained with Coomassie brilliant blue to confirm the presence of all GST fusion proteins.

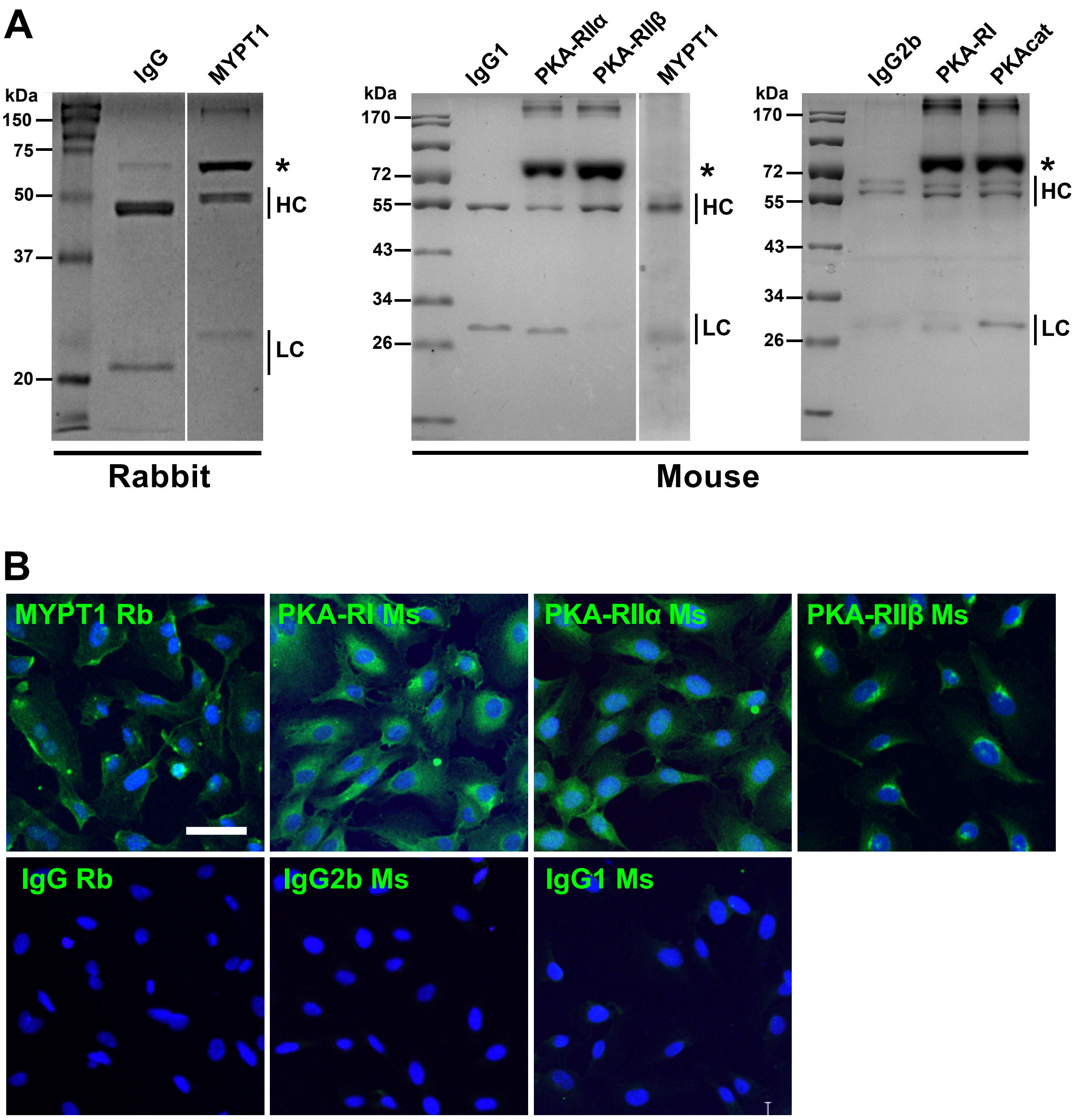


**Supplemental Figure 2. Characterization of monoclonal antibodies used for immunoprecipitation, immunofluorescence and *in situ* proximity ligation assay. (A)** Monoclonal antibodies against MYPT1 and PKA subunits and their matching immunoglobulin isotype controls were resolved by SDS-PAGE and gels were stained with Coomassie brilliant blue. 0.5 µg of antibody raised in rabbit against MYPT1 and its matching immunoglobulin control or 1 µg of antibodies raised in mouse against MYPT1 and PKA subunits and their matching immunoglobulin isotype controls were loaded. HC, heavy chain; LC, light chain; * protein (bovine serum albumin) used as stabilizer in some commercial antibodies. **(B)** Immunofluorescence staining of MYPT1 and PKA-R subunits in HUVECs**.** Cells were fixed, permeabilized and incubated with the indicated antibody followed by species-specific fluorescently labelled Alexa fluor488 conjugated secondary antibodies (green). Nuclei were counterstained with DAPI (blue). Images were acquired with a ZEISS LSM 710 confocal microscope equipped with a 20× objective. Representative images are central slices from Z stacks taken at 1.80 µm intervals. Scale bar represents 50 µm and applies to all images.

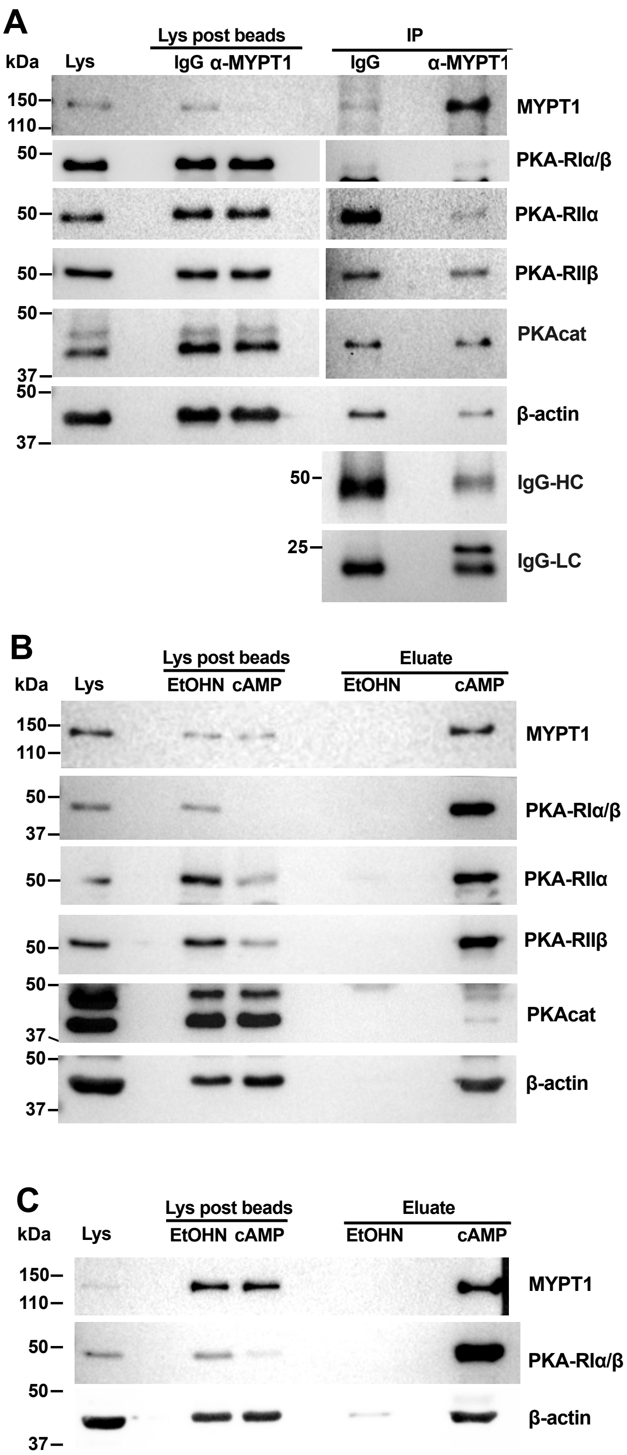


**Supplemental Figure 3. Immunoprecipitation and cAMP affinity pulldown of MYPT1 in platelets.** (**A**) Immunoprecipitation of MYPT1. Human platelet lysates were subject to immunoprecipitation with MYPT1-specific antibodies. The same isotype of the total immunoglobulin G (IgG) was used as a control. Protein complexes were examined by western immunoblot for the presence of PKA-R subunits as well as for the PKAcat subunit and for the immunoglobulin heavy chain (HC) and light chain (HC). Lys, total cell lysate. **(B)** cAMP affinity pull-down. Platelet lysates (500 µg) were incubated with 25 µl of 8-AHA-cAMP (cAMP) agarose beads or control ethanolamine (EtOHN) agarose beads. Attached complexes were examined by western immunoblot for the presence of MYPT1 and PKA subunits. **(C)** cAMP affinity pull-down in the presence of latrunculin B. Washed platelets were incubated with 20 µM latrunculin B for 30 min at 37°C, lysed and processed as in B. Beta-actin (β-actin) was used as a loading control for lysates in A, B and C. Immunoblots are representative of at least three independent experiments.

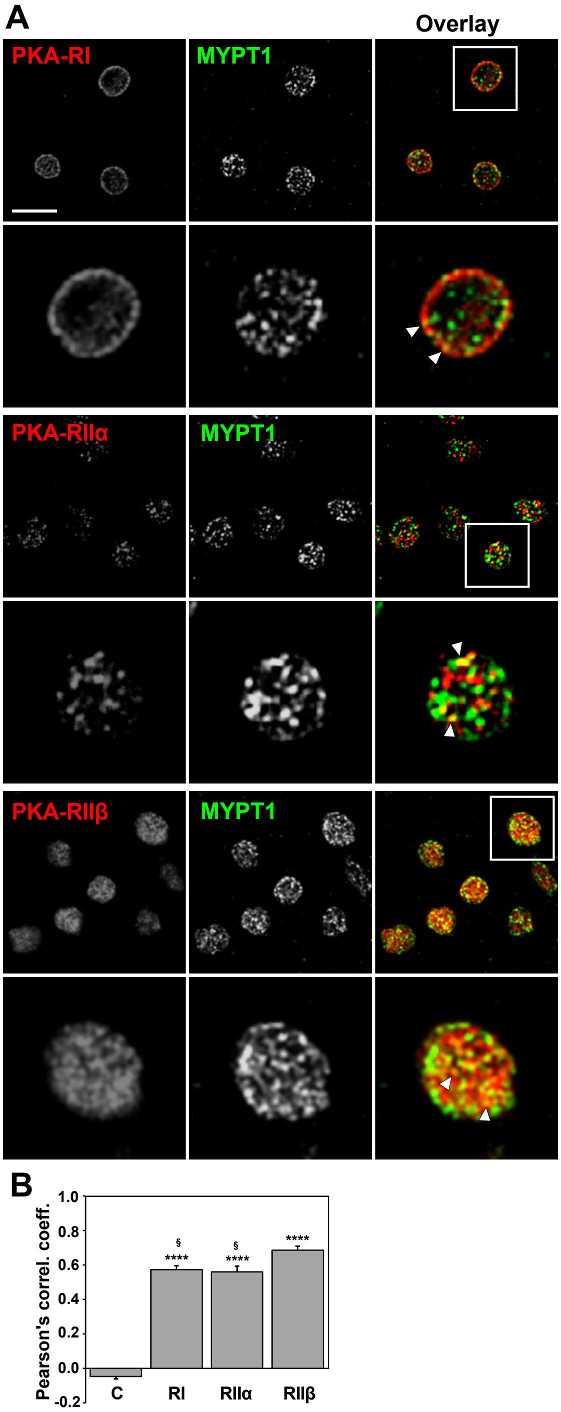


**Supplemental Figure 4. Double immunofluorescence staining of MYPT1 and PKA-R subunits in resting platelets. (A)** Human washed platelets were fixed in suspension with 4% paraformaldehyde, spun on poly-L-lysine coated coverslips, permeabilized and doubled stained with rabbit anti-MYPT1 and mouse anti-PKA-RI, RIIα or RIIβ antibodies. This was followed by species-specific fluorescently labelled Alexa fluor568 (red) and Alexa fluor488 (green) conjugated secondary antibodies. Images were acquired with a ZEISS ApoTome.2 fluorescence microscope equipped with a 63× oil immersion objective and deconvolved. Representative images are central slices from Z stacks taken at 0.15 µm intervals through the platelet. White arrowheads indicate instances of co-localization in the zoomed images. Scale bar represents 5 µm and applies to all non-zoomed images. **(B)** Pearson's correlation coefficients for MYPT1 and PKA-R subunit double stainings. For each condition, data are mean ± SEM of 20 platelets from 2-3 independent images. Images were analyzed as described in the Materials and Methods section. **** $p<0.0001$ relative to the control (C); § $p<0.05$ relative to PKA-RIIβ (Kruskal-Wallis test followed by Dunn's test).

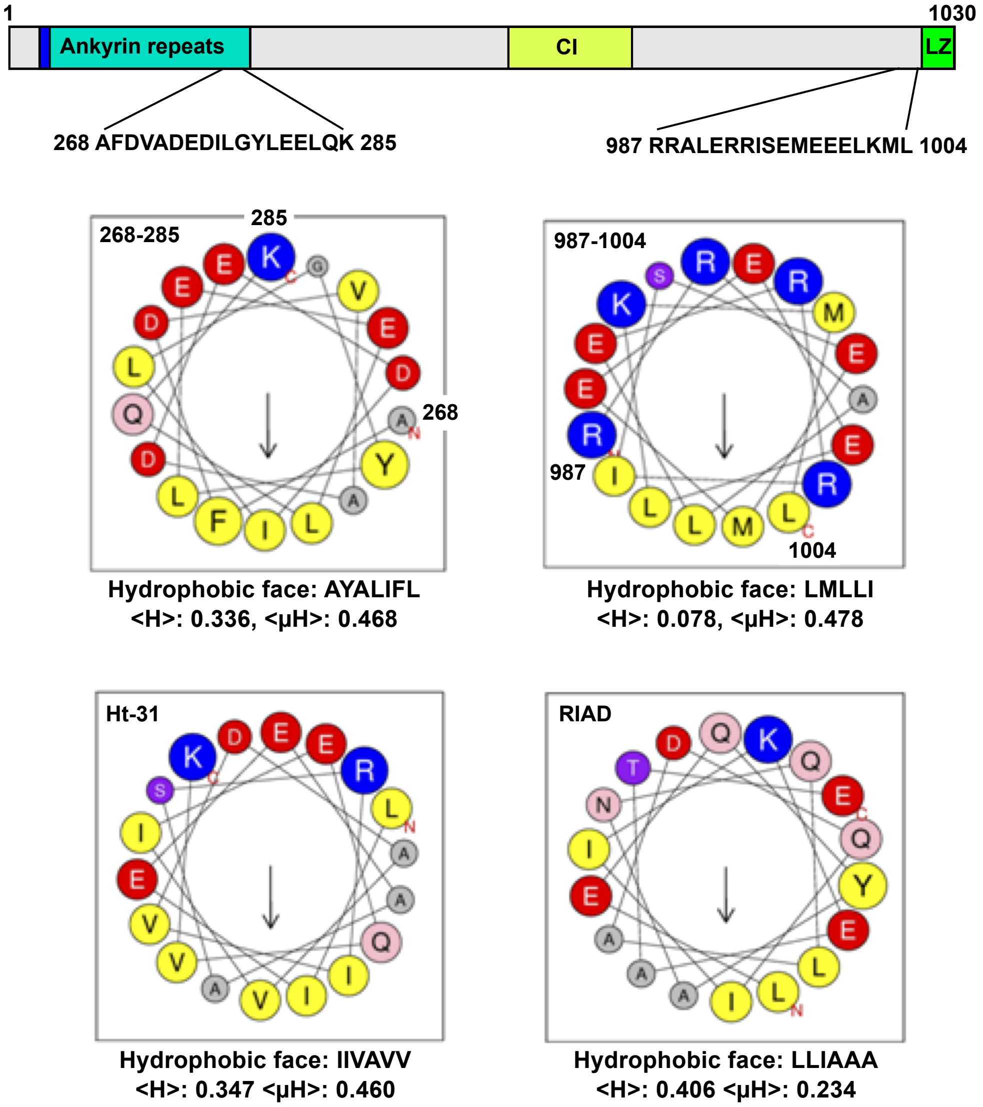


**Supplemental Figure 5. Identification of potential amphipathic helices in MYPT1.** The MYPT1 protein sequence was analyzed using the Screening α-Helix tool of the HeliQuest software (https://heliquest.ipmc.cnrs.fr). Five sequences that could potentially form amphipathic helices were further examined with the Analysis tool, reducing the candidates to two regions, 268-285 and 987-1004. Their localization in the MYPT1 sequence is indicated in the top diagram, which depicts the domain architecture of the protein: PP1c binding domain (blue), ankyrin repeats, central insert (CI) and leucine zipper (LZ) motif. Helical wheel representations of the 268-285 and 987-1004 regions are shown, along with helical wheel representations of AKAP disruptor peptides Ht-31 and RIAD. Yellow, hydrophobic residues; blue, basic residues; red, acidic residues; pink, asparagine and glutamine; green, proline; purple, serine and threonine; gray, other residues. The position of first (N) and last (C) residues are indicated. The arrow represents the direction of the hydrophobic moment. Hydrophobic moment (<μH>) and mean hydrophobicity (<H>) values are indicated for each helix.

**A**

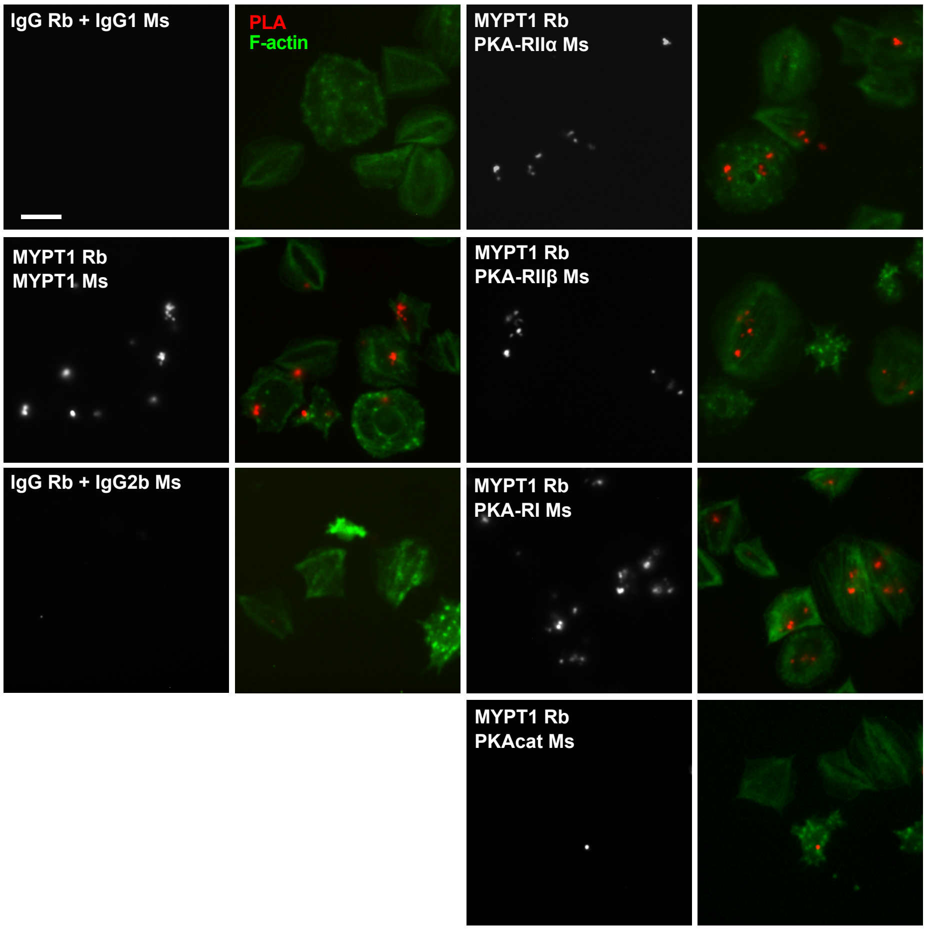


**B**

| Rabbit antibody | IgG | MYPT1 | | IgG | MYPT1 | | |
|---|---|---|---|---|---|---|---|
| Mouse antibody | IgG2b | PKAcat | PKA-RI | IgG1 | PKA-RIIα | PKA-RIIβ | MYPT1 |
| Number of dots | % Platelets | | | | | | |
| 0 | 90.1 | 81.4 | 45.7 | 91.9 | 56.0 | 63.6 | 45.9 |
| 1 | 7.4 | 8.4 | 22.8 | 5.4 | 20.9 | 12.0 | 24.1 |
| 2 | 2.5 | 6.8 | 5.5 | 1.8 | 7.6 | 8.9 | 13.2 |
| 3 | 0.0 | 1.3 | 12.3 | 0.9 | 7.1 | 5.4 | 5.3 |
| 4 | 0.0 | 1.3 | 4.1 | 0.0 | 4.9 | 3.1 | 3.0 |
| 5 | 0.0 | 0.4 | 2.7 | 0.0 | 2.2 | 3.9 | 3.8 |
| ≥6 | 0.0 | 0.4 | 6.8 | 0.0 | 1.3 | 3.1 | 4.9 |
| Mean ± SEM | 0.12 ± 0.04 | 0.37 ± 0.06 | 1.48 ± 0.14 | 0.12 ± 0.04 | 1.01 ± 0.11 | 0.97 ± 0.10 | 1.37 ±0.12 |
| % platelets with ≥1 dots | 9.9 | 18.6 | 54.3 | 8.1 | 44.0 | 36.4 | 54.1 |
| Number of platelets scored | 81 | 237 | 219 | 111 | 225 | 258 | 266 |
| Statistical significance (P) | | 0.0564 | $2.78 \times 10^{-14}$ | | $8.61 \times 10^{-10}$ | $9.14 \times 10^{-8}$ | $4.44 \times 10^{-16}$ |

**Supplemental Figure 6. *In situ* proximity ligation assay for MYPT1 and PKA-R subunits in platelets**. **(A)** Human washed platelets were allowed to spread on fibrinogen for 45 minutes, fixed, permeabilized and incubated with the indicated mouse (Ms) and rabbit (Rb) antibodies, followed by PLA reaction (red dots). The left panels are a positive control (mouse and rabbit anti MYPT1) and isoptype specific immunoglobulin negative controls (mouse IgG2b for anti-PKA-RI and IgG1 for anti-PKA-RIIα, PKA-RIIβ and MYPT1, rabbit IgG for anti-MYPT1). Cells were counterstained with FITC-phalloidin for F-actin (green). Images were acquired with a ZEISS ApoTome.2 fluorescence microscope equipped with a 63× oil immersion objective and deconvolved. Scale bar represents 5 μm. **(B)** Quantification of the *in situ* PLA assay. Dots were scored in the indicated numbers of platelets and the frequency of each class expressed as percentage. Statistical significance was calculated relative to the respective pair of isotype-specific immunoglobulins (Kruskal-Wallis test followed by Dunn's test).

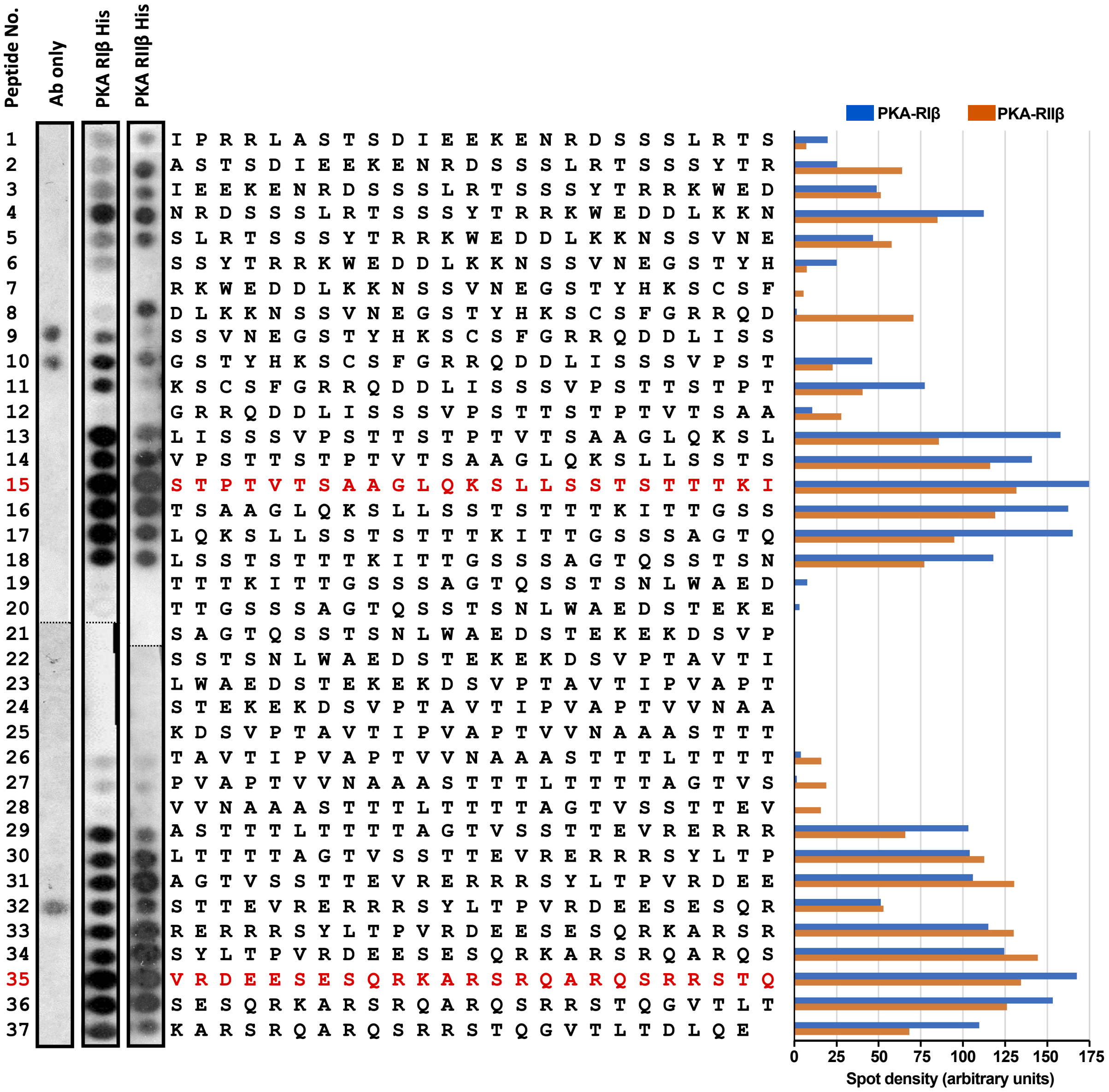


**Supplemental Figure 7. Peptide array epitope mapping of PKA-R subunits onto MYPT1.** An array of overlapping 25mers, sequentially shifted by 5 amino acids, spanning MYPT1-C2 protein segment (I501–E706) was overlaid with recombinant His-tagged PKA-RIβ or RIIβ, probed with HRP conjugated anti-polyhistidine antibody and visualized by ECL. For the antibody only control (Ab only), PKA-R was omitted. Spot densities are shown in arbitrary units after subtraction of the corresponding antibody only control density. Binding peptides selected for follow up analysis are indicated in red.

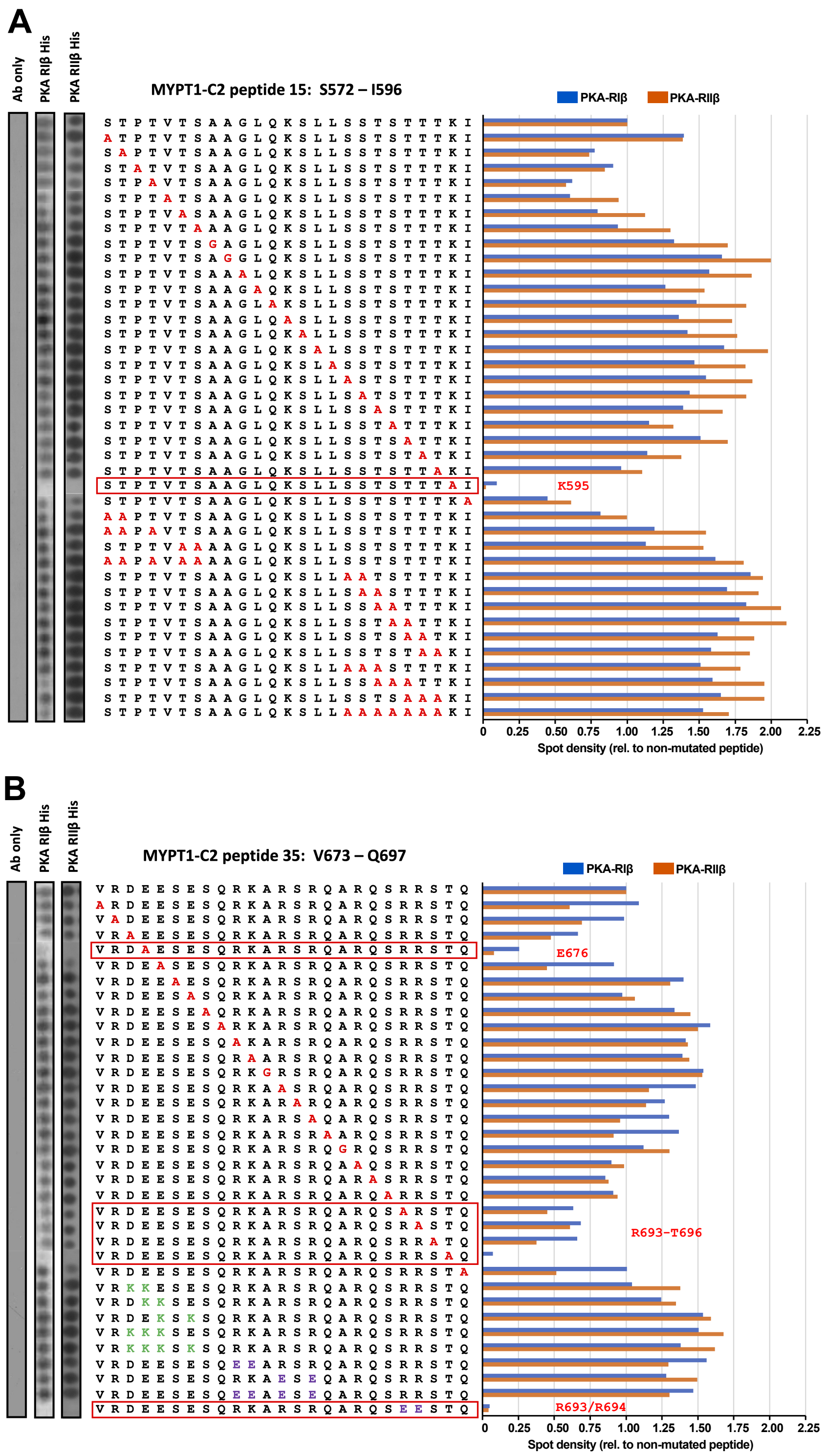


**Supplemental Figure 8. Peptide array substitution analysis of MYPT1-C2 peptides.** Arrays were overlaid with recombinant His-tagged PKA-RIβ or RIIβ, probed with HRP conjugated anti-polyhistidine antibody and visualized by ECL. For the antibody only control (Ab only), PKA-R was omitted. **(A)** Peptide array substitution analysis of MYPT1-C2 peptide 15. Key MYPT1-C2 hot spots were identified by single, dual or multiple alanine substitutions. The red box highlights a peptide where a substitution resulted in almost complete loss of binding. **(B)** Peptide array substitution analysis of MYPT1-C2 peptide 35. Key MYPT1-C2 hot spots were identified by single alanine substitutions or by opposite charge residue substitutions. The red boxes highlight peptides where substitutions resulted in consistent reduction or complete loss of binding. Spot densities are shown relative to the non-mutated peptide after subtraction of the corresponding antibody only control density.
